# Gap-free X and Y chromosomes of *Salix arbutifolia* reveal an evolutionary change from male to female heterogamety in willows, without a change in the sex-determining region

**DOI:** 10.1101/2023.10.11.561967

**Authors:** Yi Wang, Guangnan Gong, Rengang Zhang, Elvira Hörandl, Zhixiang Zhang, Deborah Charlesworth, Li He

**Author notes:** Author for correspondence. Deborah Charlesworth,; Li He.

## Abstract

In the *Vetrix* clade of *Salix*, a genus of woody flowering plants, sex determination involves chromosome 15, but an XY system has changed to a ZW system. We used genome sequencing (with chromosome conformation capture (Hi-C) and PacBio HiFi high-fidelity reads) to study the evolutionary history of the sex-linked regions before and after the transition. We assembled chromosome level gap-free X and Y chromosomes of *Salix arbutifolia*, and distinguished the haplotypes in the 15X- and 15Y-linked regions. This revealed “micro-heteromorphism” differentiating the haplotypes of the Y- and X-linked regions, including insertions, deletions and duplications. Unusually, the X-linked region is considerably larger than the corresponding Y region, and we show that this primarily reflects extensive accumulation of repetitive sequences and gene duplications. The phylogenies of single-copy orthogroups within the sex-linked regions of *S. arbutifolia* (X and Y) and *S. purpurea* (Z and W) indicate that they possess a common ancestral sex-linked region that is physically small and located in a repeat-rich region near the chromosome 15 centromere. During the change in heterogamety, the W-linked region was derived from the X-linked one and the Z from the Y. The W may subsequently have evolved a region in which recombination became suppressed. We also detected accumulation of genes with opposite sex-biases in the sex-linked regions.

## Introduction

Genetic sex determination systems have evolved independently as dioecy has evolved in many different angiosperm lineages, and both male heterogametic (XX/XY) and female heterogametic (ZZ/ZW) systems are recorded (Westergaard, 1958; Bull, 1985; Ming et al., 2011). The evolution of sex chromosomes is thought to involve several steps initiated by the evolution of a sex-determining gene or genes within a genome region. In plants, this initial step may involve mutation in a gene with an essential male function (denoted by M), creating females, followed by a femaleness-suppressing mutation (SuF) that created males and defines a Y-linked region (Charlesworth and Charlesworth, 1978). Such an incompletely Y-linked region carrying the M and SuF factors creates selection favouring closer linkage between the two genes, which may lead to complete recombination suppression, producing Y- and X-linked regions that thereafter evolve without exchanges, so that the Y becomes a differentiated sex-linked region (SLR). Such regions often accumulate repetitive elements (Charlesworth et al., 1994), directly reducing gene density compared to homologous X-linked regions, and even more markedly compared with autosomal or pseudoautosomal regions (PARs), as seen, for example, in *Silene latifolia* (Bergero et al., 2008). After enough evolutionary time, genetic degeneration may occur, eventually sometimes leading to major gene losses, creating hemizygosity of genes and regions (Charlesworth et al., 2005). The absence of recombination also allows the accumulation of Y-specific gene duplications and chromosome rearrangements and eventual changes in size and heterochromatinization of the sex chromosomes (Charlesworth et al., 2005; Bergero and Charlesworth, 2009). W chromosomes in species with female heterogamety undergo changes similar to those observed for Y chromosomes (Fraisse et al., 2017; Picard et al., 2018; Xu et al., 2019c).

In contrast, the X and Z chromosomes, which recombine in one sex, largely retain the characteristics of their progenitors, though some changes are predicted (Mrnjavac et al., 2023); for example, in XY species in which recombination occurs in both sexes, the X recombines only in females, so that its recombination rate is lower than that for the autosomes. However, empirical data concerning X chromosome changes in dioecious plants are limited, though an assembly of the *Silene latifolia* X found a lower gene density than that on most autosomes (Yue et al., 2023). The *Salix* clade *S. dunnii* has a XY system on chromosome 7, and comparison with the non-sex chromosomal homologs in related species suggests that it has an expanded X-SLR (He et al., 2021). Distinguishing between Y- and X-SLRs can detect “micro-heteromorphism” between them at the molecular level, including the predicted accumulation of transposable elements (TEs) and other repetitive sequences (see above), as well as other rearrangements between two haplotypes.

Reliable assemblies of both sex-linked regions are needed to study these processes, including to quantify the extent of genetic degeneration in dioecious angiosperms. Degeneration is known to have been initiated in some species (Bergero and Charlesworth, 2011; Chibalina and Filatov, 2011; Crowson et al., 2017; Prentout et al., 2021), but is currently not well characterized in any species. To better understand degeneration, the proportion of genes retained on a Y or W chromosome should be estimated, using reliably documented sets of X or Z-linked genes to infer the genes that were ancestrally present, as has been done in the human XY pair (Wilson Sayres and Makova, 2013), but in few other species, even among animals.

Reliable assemblies of both sex-linked regions are also needed for determining the sizes of the sex-linked regions, and for testing whether Y (or W) regions have undergone recombination suppression, and, if so, to study the mechanism(s). Although, as described above, recombination suppression may be initiated by the emergence of the sex-determining genes, in some species it has spread along the sex chromosome, forming “evolutionary strata” with different levels of Y-X sequence divergence, reflecting different times when recombination stopped (Lahn and Page, 1999). Data on evolutionary strata are scarce, even for animals. Y-X divergence for synonymous sites is lower for genes close to the X-PAR than for ones more distant from the PAR, in the flowering plants *Silene latifolia* and in the related species *Cannabis sativa* and *Humulus lupulus*, all of which have large completely Y-linked regions (Lahn and Page, 1999; Papadopulos et al., 2015; Prentout et al., 2021). Lahn and Page (1999) suggested that such strata reflect fixation of inversions on sex chromosomes, and inversions were found in the papaya sex-linked region (Wang et al., 2012). Flowering plants are well-suited to testing for inversions, because dioecy has often evolved recently, so that there has not been enough time for later rearrangements to obscure the events involved in the early stages. However, long-read sequence data from which they might be identified, if present, are not yet available for many dioecious plants, and data from recently evolved sex-determining regions are scanty.

The Salicaceae are therefore an interesting family, as turnovers have occurred, creating young sex-determining regions. Strata may have formed in the SLR of *Populus euphratica*, and an inversion detected in the assembly could have been involved, as the estimated mean divergence for genes within the inversion was 4%, versus only 0.018 for the rest of the SLR (Zhang et al., 2022b); however definitive evidence has not yet been published. In *Salix viminalis*, two evolutionary strata were identified, but the assemblies were low-quality, especially in the sex-linked region; moreover, the authors concluded that recombination may not be completely suppressed (Almeida et al., 2020). Further studies in this family are therefore needed.

Sex chromosome turnovers are classified into homogametic (XY to XY, or ZW to ZW) and heterogametic transitions (XY to ZW, or ZW to XY) (Bull, 1983; Saunders et al., 2018), and (van Doorn and Kirkpatrick, 2010) introduced the categories of nonhomologous transitions (the sex-linked region is on a different chromosome, and the old sex chromosome or linkage group becomes an autosome) versus homologous transitions (with no major change in location of the sex determining region); predictions for these two types of changes were reviewed by (Bull, 1983). A change to a different region within the same chromosome is also possible (Ieda et al., 2018; Kabir et al., 2022). The Salicaceae display all these types (Yang et al., 2021; Li et al., 2022; Wang et al., 2023). It is hypothesized that, in heterogametic transitions, W-linked regions are likely to derive from ancestral X-linked regions (or vice versa), because these carry feminizing genes, while the Z and Y may often carry masculinizing genes (Miura, 2007; Ma et al., 2018; Ogata et al., 2018; Furman et al., 2020).

*Salix* includes two main clades: the *Salix* and *Vetrix* (Gulyaev et al., 2022). In the *Vetrix* clade, an XY to ZW transition has been found, with both having physically small sex-linked regions being on chromosome 15 (Wang et al., 2022; Wang et al., 2023). The extant species with female heterogamety, which we abbreviate to 15ZW (e.g., *S. purpurea*, *S. viminalis*, and *S. polyclona*) and *S. arbutifolia* (with male heterogamety, or 15XY) share an ancestral 15XY system, but high-quality X and Y reference assemblies of *S. arbutifolia* are currently lacking, which hinders understanding of these evolutionary changes. (Hu et al., 2022) attempted to use another 15XY species, *S. exigua*, to study the XY to ZW transition. However, phylogenetic incongruence between *S. exigua* in chloroplast and nuclear trees implied complicated origination of this species (Gulyaev et al., 2022).

Here, we asked whether sex-linked regions in these species carry the same genes, using *Salix arbutifolia*, which is at the key phylogenetic position for inferences about this the change in heterogamety. As the sex-linked contigs of the previous reference genome of *S. arbutifolia* (Wang et al., 2022) are low quality, we used chromosome conformation capture (Hi-C) and PacBio HiFi (high-fidelity) reads to assemble chromosome level gap-free X and Y chromosomes. A first aim was to ask whether a cytokinin response regulator gene, ARR17-like gene is involved in these species, as in two other Salicaceae species, *Populus deltoides* and *P. tremula*, both with male heterogamety; in both, partial *ARR17*-like gene duplicates give rise to small RNAs apparently causing male-specific DNA methylation and silencing of the ARR17-like gene. A sex-determining function was confirmed by CRISPR-Cas9-induced mutation, and it has been proposed that such a gene could be involved in sex-determination across the Salicaceae (Muller et al., 2020; Xue et al., 2020; Yang et al., 2021). When the ARR17-like gene is expressed, the individual develops into a female, and when it is silenced or absent, it develops into a male. If such a gene is also involved in sex-determination in the XY and/or ZW *Salix* systems studied here, they should be located in the oldest-established sex-linked region(s). We also: (ii) investigate the organization of the X- and Y-linked regions, and (iii) compare the evolution of the 15XY *S. arbutifolia* and 15ZW *S. purpurea* sex-linked regions since their divergence.

## Results

### Genome assembly

In brief, our haplotype-resolved assembly of the *S. arbutifolia* genome was derived by integrating a total of 45.2 Gb Illumina reads, 44.9 Gb of HiFi reads with an average length of 18 kb, and 44.7 Gb of Hi-C reads (Table S1). A 642.58 Mb primary assembly with 353 contigs was obtained using Hifiasm from the HiFi reads; its contig N50 was 13.47 Mb. The Hi-C reads were then used to scaffold the contigs to a chromosome-scale assembly, resulting in 40 scaffolds. 38 of these were assigned to the 19 *Salix* chromosomes, based on synteny with *S. brachista*. Most assembled chromosomes were gap free (except for chr04a, chr07a, chr07b and chr19a). The mitochondrial and chloroplast genomes formed two other scaffolds, of 659,371 and 155,219 bp, respectively. The final nuclear chromosomal assembly was 612 Mb, with a contig N50 of 16 Mb and scaffold N50 of 16.45 Mb (Table 1), including two complete haplotypes of each region.

**Table 1:**
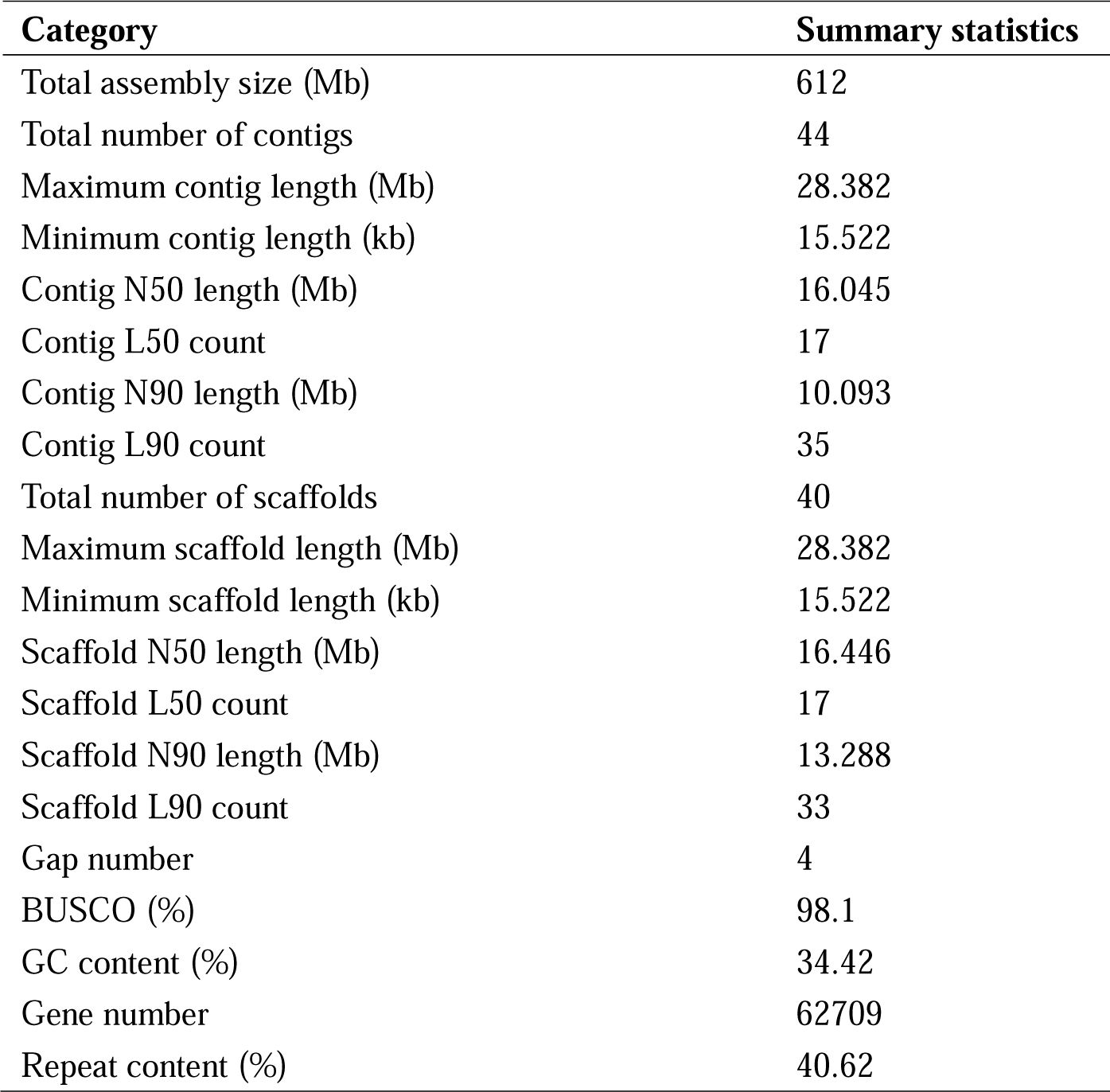
Statistics of the *Salix arbutifolia* genome assembly.

| Category | Summary statistics |
| --- | --- |
| Total assembly size (Mb) | 612 |
| Total number of contigs | 44 |
| Maximum contig length (Mb) | 28.382 |
| Minimum contig length (kb) | 15.522 |
| Contig N50 length (Mb) | 16.045 |
| Contig L50 count | 17 |
| Contig N90 length (Mb) | 10.093 |
| Contig L90 count | 35 |
| Total number of scaffolds | 40 |
| Maximum scaffold length (Mb) | 28.382 |
| Minimum scaffold length (kb) | 15.522 |
| Scaffold N50 length (Mb) | 16.446 |
| Scaffold L50 count | 17 |
| Scaffold N90 length (Mb) | 13.288 |
| Scaffold L90 count | 33 |
| Gap number | 4 |
| BUSCO (%) | 98.1 |
| GC content (%) | 34.42 |
| Gene number | 62709 |
| Repeat content (%) | 40.62 |

About 99.6% of the Illumina short reads mapped to the genome assembly, and about 99.4% of the assembly was covered by at least 20× reads. About 99.6% of the HiFi reads mapped to the genome assembly and 99.4% were had coverage of at least 20 (Table S2). BUSCO analysis showed that the primary assembly covered 95.9% of complete BUSCO genes. A further 0.7% had fragmented matches to other conserved genes, and 3.4% were missing. Thus, our genome assembly has high continuity, coverage, and accuracy.

### Genome Annotation

A total of 937,169 repetitive sequences were identified, accounting for an estimated 40.62% of the assembled genome (or a total of 248.46 Mb). Among TEs, LTR-RTs were the most abundant, accounting for 16.16% of the genome, with Gypsy and Copia class I retrotransposons (RT) accounting for 5.83% and 6.68% of the genome, respectively (Table S3). We annotated 3,793 full-length LTR retrotransposons (LTR-RTs), including 122 in the 15X-SLR and 78 in the 15Y-SLR. Divergence estimates of the intact LTR-RTs in the 15X-SLR range from 0 to 0.14 (mean 0.04, suggesting insertion times from 0 to 28.46 Mya) and 0 to 0.18 for the 15Y-linked region (mean 0.04, suggesting insertion times from 0 to 36.78 Mya) (Table S4).

Based on the transcript dataset, homology searches, and *ab initio* prediction, we identified a total of 62,709 gene models, including 58,688 protein-coding genes, 1226 transfer RNAs (tRNAs), 1383 ribosomal RNAs (rRNA) and 1412 unclassifiable noncoding RNAs (ncRNAs) in the two haplotypes (Table S5). The average *S. arbutifolia* gene is 3457.5 bp long and contains 6.2 exons (Table S6). 98.32% of the predicted protein-coding genes matched a predicted protein in public database(s) (Table S7).

### Identification of sex-linked regions of *S. arbutifolia*

The mapping ratio of the 41 wild individuals’ clean reads using our haplotype *a* and *b* genome sequences as references ranged from 86.00 to 93.80% (mean 90.55%, Table S8) and 87.40 to 94.00% (mean 91.31%, Table S9), respectively. We obtained 3,308,616 and 3,324,319 high-quality SNPs using the *a* and *b* haplotype genome sequences as references in 41 individuals of *S. arbutifolia*, respectively. This revealed a chromosome 15 region with chromosome quotient (CQ) close to 2 on 15*a*; because the value expected for autosomal sequences is close to 1, this indicates that the 15*a* haplotype in this region corresponds to 15X (16.17 Mb). Supporting this conclusion, the syntenic region has a CQ close to 0 in the 15*b* haplotype, consistent with it representing the 15Y (10.98 Mb) (Fig. S1; Fig. S2). Changepoint analysis of F_ST_ between male and female individuals detected high values in a 7.06 Mb region located between 3.64 and 10.7 Mb of 15X, compared with the flanking regions, confirming that this is the 15X-SLR. The regions located at both ends of the chromosome assembly will therefore be termed X-PAR1 and X-PAR2; their total length is 9.11 Mb (Fig. 2a; Fig. S3). The 15Y-SLR signal is identified in a smaller region, of 1.81 Mb, between 3.62-5.43 Mb on 15Y, while the total length of the Y-PARs is 9.17 Mb, similar to the X value, as expected (Fig. 2e; Fig. S4).

In the X- and Y-SLRs, our GWAS detected 34 and 274 SNPs (Table S10) significantly associated with sex (Fig. 2a, e; Fig. S5; Fig. S6). These SNPs identify one sex-linked gene in the X-SLR and seven in the Y-SLR. Among them, the Saarb15bG0044200 reciprocal best hit in *A. thaliana* is BRR2C, a gene encoding the DExH-box ATP-dependent RNA helicase, a highly conserved spliceosome component required for efficient splicing of flowering locus c (FLC) introns in this species, as well as regulation of flowering locus t (FT) and suppressor of overexpression of constans 1 (SOC1) (Mahrez et al., 2016). Another sex-linked gene with an *A. thaliana* hit is Saarb15bG0048300, which may be the ortholog of the PPRT3 gene, which plays a regulatory role in the abscisic acid (ABA) signaling pathway. Table S10 gives detailed information about the other significant sex-linked genes, which had no evident likely reproductive functions. Homology searches (see Methods) identified 1 and 3 pseudogenes in 15X-SLR and 15Y-SLR, respectively (Table S11).

### Micro-heteromorphism of X- and Y-SLRs of *S. arbutifolia*

We detected lower gene density (and higher repeat density) in the X- and Y-SLRs than in the PARs (Table 2), suggesting repetitive element accumulation in these regions. The X-SLR repeat sequence density is extremely high, at 80.73%, much higher even than in the Y-SLR (64.09%; Table 2). The repeat sequence density in the two PARs is only 30.86%. The X- and Y-SLRs both have more LTR-Gypsy and LTR-Copia retrotransposons than the PARs (Table 2, Fig. 2b, f).

**Table 2:**
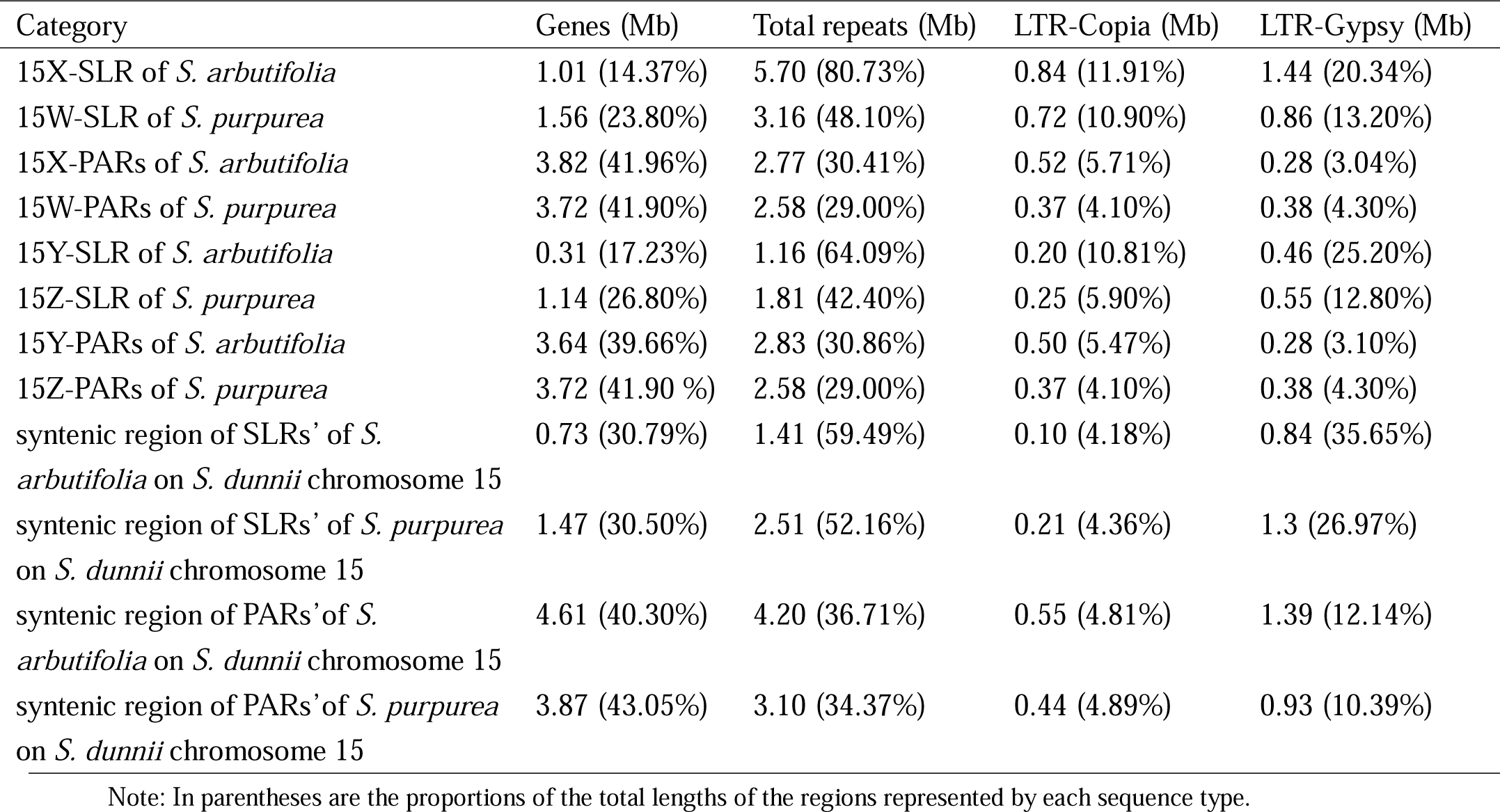
Size of genes and LTR retrotransposons in different regions of the genome. In parentheses are the proportions of the total lengths of the regions represented by each sequence type.

Synteny analysis detected a ∼4.36 Mb region (4.98-9.34 Mb) specific to the X-SLR (Fig. 1b), which we term the “extended region”. 86.50% of this region consists of repetitive sequences, 38.22% of which are unknown repeat types, 18.08% are LTR-Gypsy sequences, and DNA-Helitron and LTR-Copia formed 13.50% and 11.21%, respectively (Table S12).

**Figure 1:**
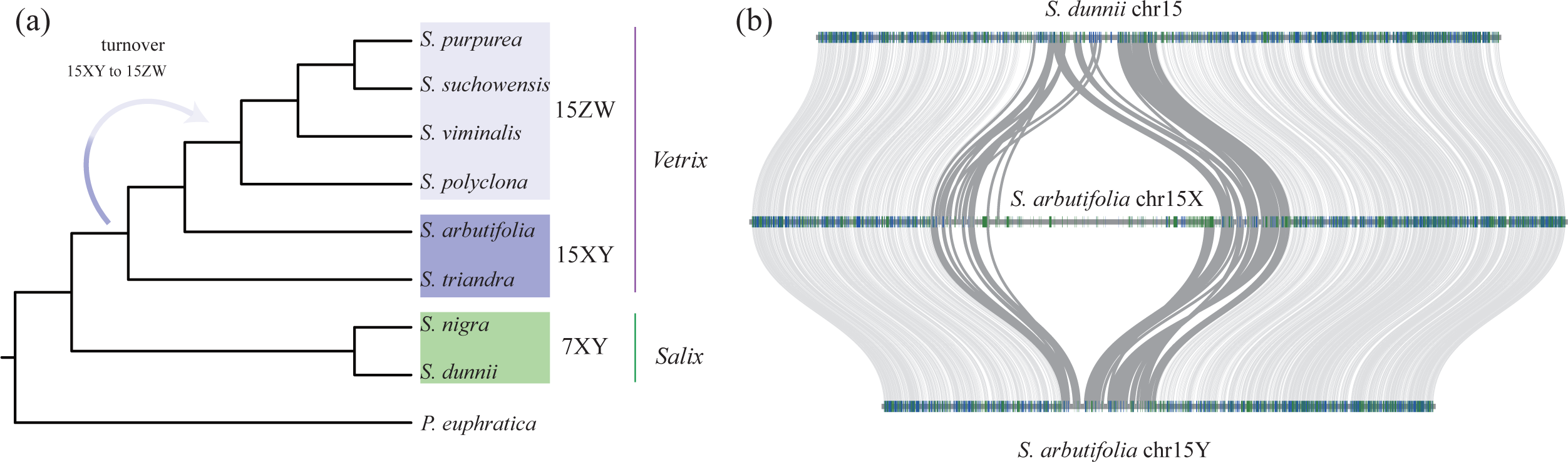
(a) Phylogenetic relationships of species with known sex determination systems in *Salix* and turnover events in the *Vetrix* clade, adapted from Wang, et al., (2023). (b) Microsynteny between the *Salix dunnii* autosome 15 and the *Salix arbutifolia* sex chromosomes 15X and 15Y, showing the micro-heteromorphism due to a large region present only in the 15X sex-linked region. The grey lines are inferred to be within the *S. arbutifolia* sex-linked regions, as well as the corresponding *S. dunnii* region.

Within the sex-linked regions we found many duplicated genes, especially in the X-SLR: 174 (60.21%) of the X-SLR genes are duplicates, and 19 (19.2%) of the Y-SLR genes. Of the duplicated genes in the 15X-SLR, 134 are in the “extended region” and occupy a total length of 0.51 Mb (11.67% of the region) (Fig. 2d, Table S13). Together, the repeats and these duplicated genes constitute 98.17% of the extended region, which thus lacks single-copy genes. Interestingly, this X-linked region includes a tandem array of 34 homologs of the 15Y-SLR Saarb15bG0044200 gene (Table S14).

**Figure 2:**
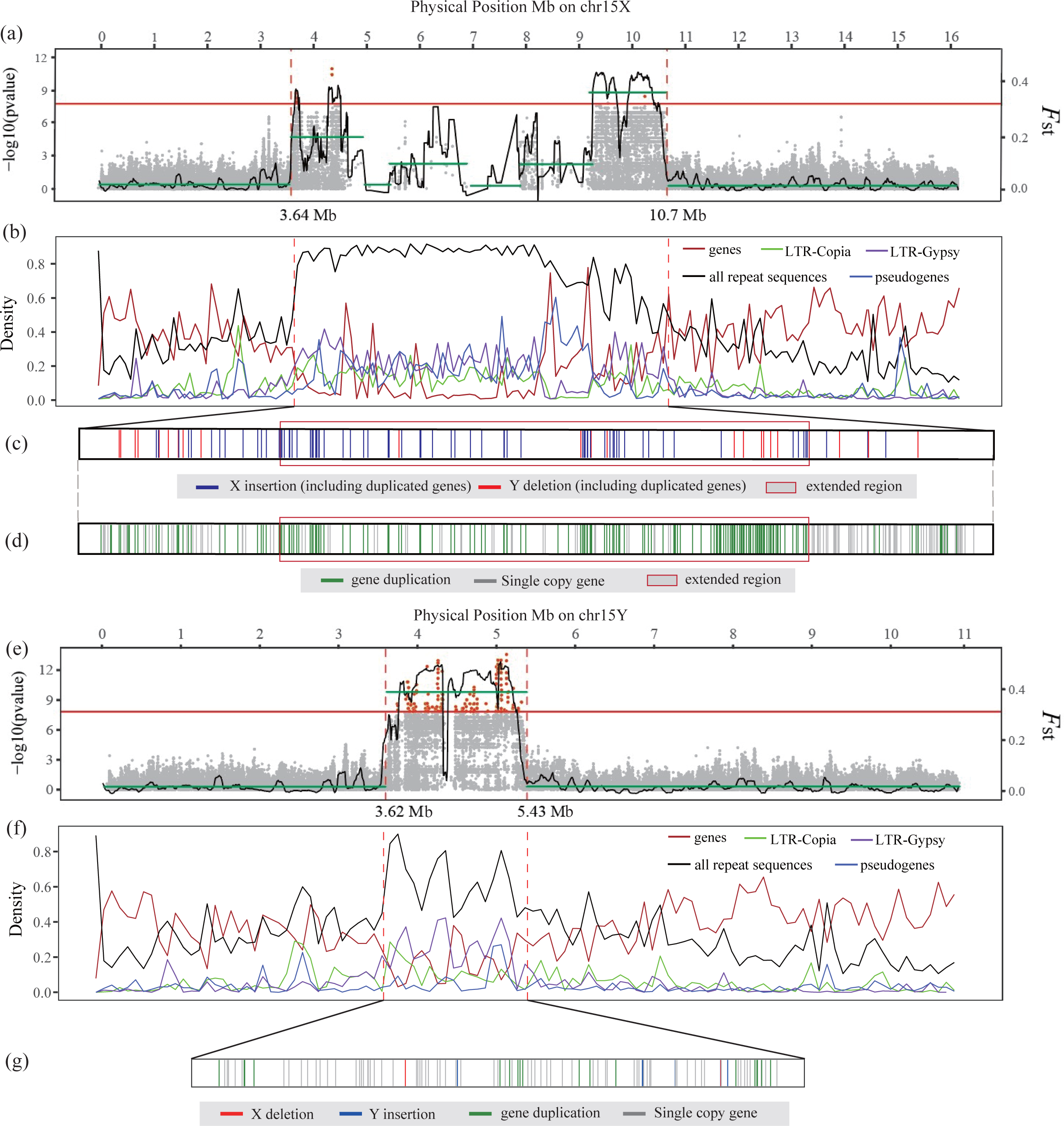
Identification of the *S. arbutifolia* 15X and 15 Y-SLRs. a-d, Analysis of the 15X region. (a) Gray dots represent *p*-values for all 15X SNPs analyzed by GWAS, and red dots above the red line indicating the Bonferroni-corrected significance level corresponding to α < 0.05. The black broken line represents F_ST_ values between the sexes in 100-kb overlapping windows, with 10-kb steps. Green horizontal lines indicate three regions that changepoint analysis suggests have different F_ST_ values. (b) Densities of LTR-Gypsy (purple line), LTR-Copia (green line), all repeat sequences (black line), pseudogenes (blue line), as well as genes (red line) in the entire 15X. (c) X-SLR-specific genes (including duplicates). The blue vertical lines represent “X insertion” genes, red ones are “Y deletion” genes, and the thin red box (also in part d) indicate the “X-extended region”. (d) Gene duplication events in X-SLR. The green vertical lines indicate gene duplicates, and grey ones indicate single-copy genes. e-g, Analysis of the 15Y region. The explanations above apply to these plots, except that there is no red boxed region (no part corresponding to part c above), and, in part (g) we show Y-SLR-specific genes, with red vertical lines indicating “X deletion”, and blue ones “Y insertion” genes.

Given the high repeat content, and the differences in genome arrangements, we examined gene and TE densities across all the chromosomes, to predicted the positions of the centromeres in this species. This suggested that the SLRs are within centromeric or pericentromeric regions, consistent with their low gene densities (Fig. S7).

Comparison with the Y-SLR (see the Methods) detected 112 X-SLR specific complete genes, of which 38 (including duplicates, 26 in extended region) are in the “Y deletion” category defined using homologous genome regions in outgroup species, as described in the Methods section; intriguingly, the other 74 are “X insertion” genes (again including duplicates, 59 in extended region) (Fig. 2c, Table S15). We detected 44 XY gametolog pairs of complete single-copy genes, versus only ten Y-SLR specific genes, including two “X deletion” and eight “Y insertion” genes (Fig. 2g, Table S16).

### Changed heterogametic sex in the *Vetrix* clade

We compared the 112 *S. arbutifolia* X-specific and 10 Y-specific genes with genes in the Z- and W-SLRs of *S. purpurea*. Including duplicates, we found homologs of 25 X-SLR specific genes (Table S15) in the *S. purpurea* 15W-SLR, but not the 15Z-SLR, suggesting that these genes are retained in both X and W, but not in Y and Z (Table 3). We also detected homologs of two 15Y-SLR specific genes in the 15Z-SLR, but not in the 15W-SLR (Table 3, Fig. 3a, Table S16). These results suggest that the W originated from an ancestral X chromosome, and the Z from a Y. However, including duplicated genes, 58 genes are 15X-SLR specific and have no homologs in either the 15Z- or 15W-SLR (Table 3, Table S15). Furthermore, homologs of 15X-specific genes are not restricted to the 15W-SLR, nor are 15Y-specific genes restricted to the 15Z-SLR (Table S15 and S16). In the *S. arbutifolia* 15Y-SLR and *S. purpurea* 15Z-SLR, we found seven and eight partial ARR17-like gene duplicates, respectively, and these cluster together in the IQ tree (Fig. S8), suggesting common ancestry. Among the 15W-linked ARR17-like genes (in sex-linked region), four are intact, and are very similar to those on chromosome 19 of *S. purpurea* (Fig. S8), suggesting they were translocated to 15W after splitting from *S. arbutifolia*.

**Figure 3:**
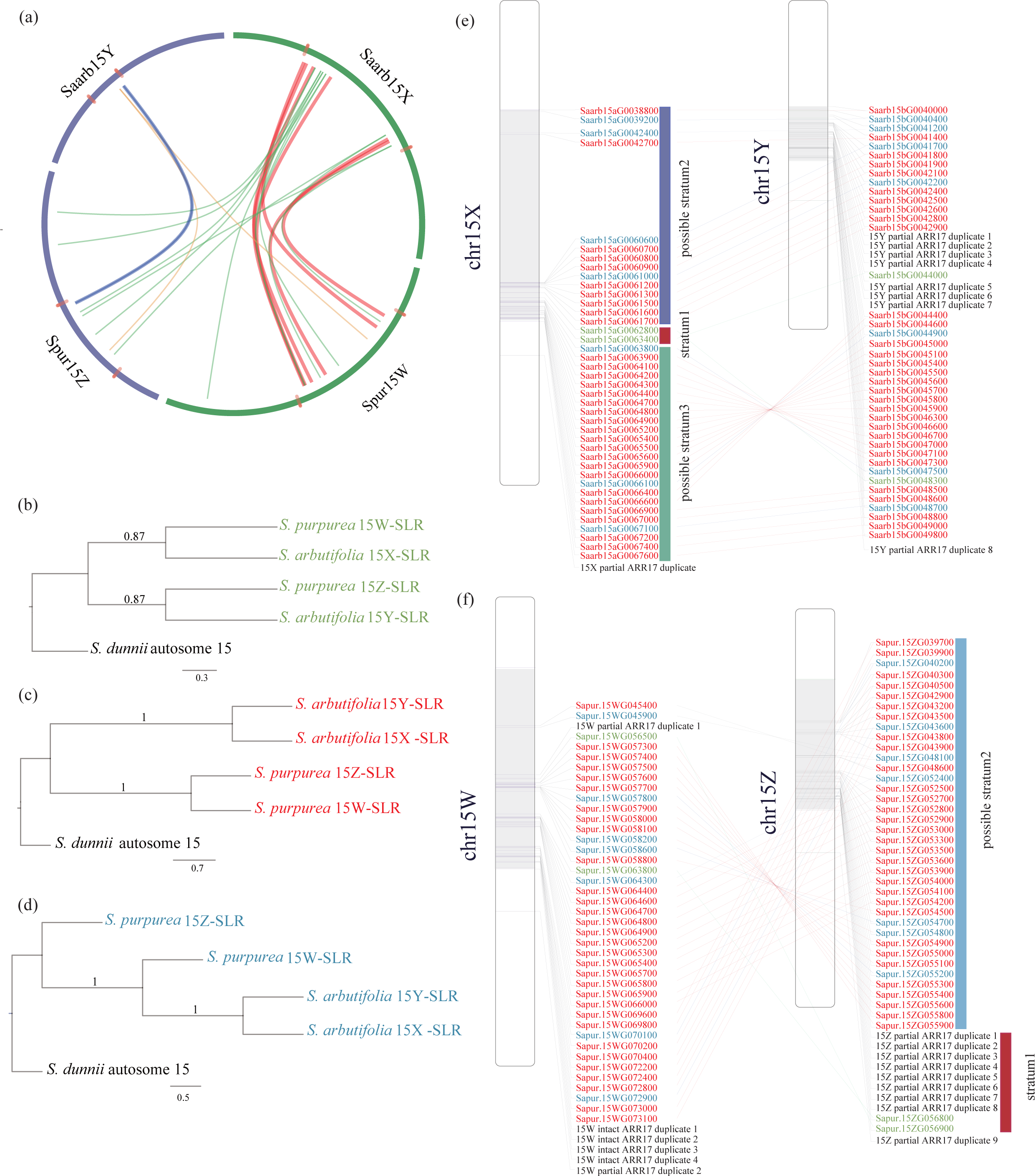
Heterogametic sex transition history of 15X, 15Y, 15Z and 15W of willows. (a) Circular plot showing *S. arbutifolia* 15X- and Y-specific genes that are present in the *S. purpurea* W- and/or Z-SLRs. The regions between the short red lines on the curved blue and green lines (chromosomes) indicate the putative sex-linked regions. Long red lines are homologous gene pairs of the 15X-SLR (specific genes) and the *S. purpurea* W-SLR; 15X-specific genes were found on both the Z and W-SLRs of *S. purpurea* are indicated by green lines. The blue and orange lines show results for S*. arbutifolia* 15Y-specific genes; homologous genes were found only on the *S. purpurea* Z-SLR are shown by blue lines, and ones on both the Z and W-SLRs by orange lines. (b) Topology I, proto-X-Y recombination stopped before the split of the two species. (c) Topology II, X-Y and Z-W recombination might stop after the split. (d) Topology III showed that 15W- and 15X and 15Y-SLRs formed a clade and sister to 15Z-SLR. (e-f) showed the positions of these single-copy genes with the three tree topologies on 15X, Y, W, and Z. The gene IDs are in colors as in parts b-d, but partial ARR17-like gene duplicates are indicated by black font. (e) 15X and Y single-copy genes, (f) 15W and Z single-copy genes. The gray boxes represent sex-linked region.

**Table 3:** The evolution relationship between specific genes on 15X and 15Y-SLRs of *S. arbutifolia* and 15Z and W-SLRs of *S. purpurea*.

| *S. arbutifolia* 15X-specific genes | X insertion<br>(including duplicated<br>genes) | Y deletion<br>(including duplicated<br>genes) | Total number |
| --- | --- | --- | --- |
| Present in the 15X and 15W-SLRs | 12 | 13 | 25 (22.32%) |
| Present in the 15X, 15W and<br>15Z-SLRs | 26 | 3 | 29 (25.89%) |
| Only present in the 15X-SLR, not<br>15ZW-SLRs | 36 | 22 | 58 (51.79%) |
| *S. arbutifolia* 15Y-specific genes | Y insertion | X deletion | Total number |
| Present in the 15Y and 15Z-SLRs | 1 | 1 | 2 (20%) |
| Present in the 15Y, 15Z and<br>15W-SLRs | 1 | 0 | 1 (10%) |
| Only present in the 15Y-SLR, not<br>15ZW-SLRs | 6 | 1 | 7 (70%) |

Among peptide sequences encoded by the SLRs of *S. arbutifolia* and *S. purpurea* and also by chromosome 15 of *S. dunnii*, we found 44 single-copy genes. Using *S. dunnii* sequences to root the trees, we found three major topologies: (I) for two single-copy genes, the X and W-linked sequences of *S. arbutifolia* and *S. purpurea* cluster together, while their Y and Z-linked genes form a well-supported separate clade, consistent with a lack of recombination before these species split (Fig. 3b) and origination of W-linked sequences from X-linked ones, and Z from Y ones; (II) for 31 single-copy genes (outside the “extended region” described above), however, the *S. arbutifolia* 15X-SLR and 15Y-SLR form one cluster, and *S. purpurea* 15Z-SLR and 15W-SLR another, consistent with the species tree, leaving it unclear whether or not recombination became suppressed (in either or both species) after the species split (Fig. 3c); (III) finally, for seven single-copy genes, the results are difficult to interpret, as the W, X and Y-SLRs formed one clade, while the 15Z-SLR sequences are sister to this clade (Fig. 3d). Possibly single-copy genes have been lost from the 15Y-SLR, and/or genes have moved from the 15X-SLR to the 15Y-SLR by ectopic or non-allelic homologous recombination (NAHR), as has been detected in human sex chromosomes (Rozen et al., 2003; Mensah et al., 2014).

### Tests for evolutionary strata on the sex chromosomes and different evolutionary trajectories in the *Vetrix* clade

Our tree topologies suggest that the fully sex-linked regions of *S. arbutifolia* and *S. purpurea* include an old evolutionary stratum with two genes that have been fully sex-linked since before the species split. We named these strata 1 in both species. It occupies 180 kb in the 15X-SLR (9.72 Mb-9.90 Mb), and 0.46 Mb in the 15Z-SLR (6.25 Mb-6.70 Mb). The partial ARR17-like gene duplicates in *S. arbutifolia* and *S. purpurea* are found near the two single-copy genes with topology I (green in Fig. 3b, e and f), or likely physically small oldest evolutionary stratum inherited from their common ancestors. The single-copy genes corresponding to topology II might reflect SLRs that emerged following the species differentiation, or younger evolutionary strata that evolved, perhaps independently in the *S. arbutifolia* 15X-SLR (possible stratum 2 and 3) and the *S. purpurea* 15Z-SLR (possible stratum 2) (see above).

As a further test for strata, we estimated Ks values for all homologous gene pairs in the possible evolutionary stratum regions, not just single-copy genes (Fig. 4b, c, S9, Table S17 and S18). In the XY species, *S. arbutifolia*, we detected significant Ks differences between evolutionary stratum 1 and possible strata 2 and 3, and the Y and X haplotypes differ by an inversion within one possible stratum 3 (Fig. 1b; Fig. 3e).

**Figure 4:**
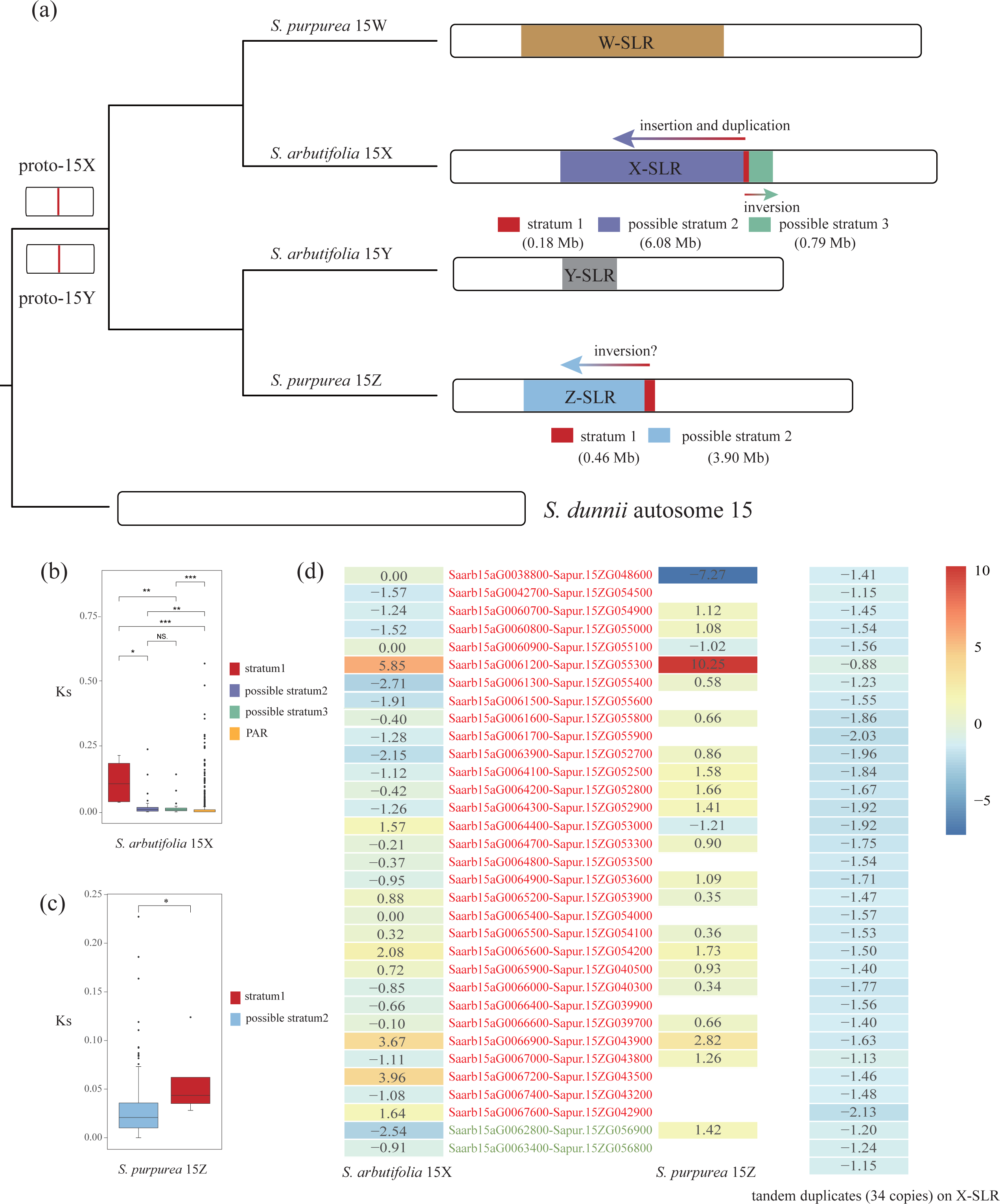
(a) Hypothetical evolutionary relationship of 15X, Y, Z, and W, and possible evolutionary strata on them. (b-c) Boxplots of Ks of all X-Y and Z-W pairs in different possible strata inferred in the *Salix arbutifolia* 15X and *Salix purpurea* 15Z, respectively (see also Fig. S9, Table S17 and S18). NS (non-significant differences) and asterisks (\**p* < 0.05, \*\**p* < 0.01, \*\*\**p* < 0.001) represent Wilcoxon tests of significance of the differences between the values of different stratum. (d) The gene expression patterns for all 33 single-copy genes with topologies I and II (left two columns), and for tandem duplicates (34 copies in total) on 15X-SLR (right column). Blank areas represent the absence of expression data.

However, this has not been shown to be male-specific, and it could reflect an inversion within a rarely recombining pericentromeric region, rather than having caused suppression of recombination for the Y-linked region. This inversion has moved the old stratum 1 single-copy gene (Saarb15bG0048300) into the possible younger stratum 3 (Fig. 3e). The values for individual genes vary considerably, and they show no clear pattern of increased values with distance from the PAR boundaries. If rearrangements occur after strata formed, they could obscure pattern of Ks values, and indeed no clear pattern is seen in the SLR of either species. The existence of younger strata remains uncertain at present (Fig. S9).

The *S. purpurea* Z-W 182 possible stratum 2 pairs have significantly (Wilcoxon tests) lower Ks values than the stratum 1 pairs (Fig. 4c, Fig. S9b, Table S18), further suggesting some or all of genes may possibly have formed younger stratum 2. Several possible inversions were identified between the *S. purpurea* Z-W chromosome pair, and could have been involved in possible stratum 2 formation (Fig. S10), or could reflect mis-assembly of SLR contigs, which relied on a linkage map to scaffold the contigs in the reference genome, and was not based on PacBio HiFi sequences (Zhou et al., 2020; Hyden et al., 2023).

Although the 15Y and 15Z regions appear to share a common ancestor, they differ in size. The *S. arbutifolia* 15Y is 10.98 Mb versus 13.25 Mb for the *S. purpurea* 15Z (Fig. 3a; Fig. 4a). Synteny analysis using the 15Z, 15Y, and *S. dunnii* autosome 15 (Fig. S11) revealed that the SLRs of both *S. arbutifolia* 15Y and *S. purpurea* 15Z have changed in size following the species split (1.81 Mb of the 15Y-SLR, but 4.36 Mb of the 15Z-SLR) (Fig. S11). Taking 15Z-SLR and the corresponding 15Y syntenic region as the likely ancestral state, the *S. arbutifolia* PAR may also have been reduced by about 1.1 Mb (Table S19); this would be consistent with formation of new strata in this species, but, in in the absence of sequences from multiple males and females, it is currently tentative. More haplotype-resolved genome sequences of willows with 15ZW and 15XY are needed to further test for strata.

### Differential expression patterns of genes in *S. arbutifolia* and *S. purpurea*

After quality control and trimming, more than 90% of our RNAseq reads from male and female catkins of B649AF and B649V mapped uniquely to the genome assembly across all samples (Table S20). A criterion of log2 M/F < −1 or >1, exclude 15X and 15Y specific genes, identified 207 female-biased genes in *S. arbutifolia* 15X, of which 54 are in the X-SLR, and 256 male-biased genes in *S. arbutifolia* 15X (19 in the X-SLR), while 455 genes on 15X were unbiased (29 in the X-SLR). Similarly, we also identified 139 female-biased genes in the 15Y assembly (three in the Y-SLR), 286 male-biased genes (37 in the Y-SLR), and 452 unbiased genes, 20 of them in the Y-SLR (Table S21). Overall, most sex-biased genes in the X-SLR are female-biased, whereas most in the Y-SLR display male-biased expression. The tandem array of 34 genes within the 15X-SLR all showed female-biased expression (Fig. 4d; Table S22). Among the 33 15X-SLR single-copy genes with topologies I or II, 12 showed female-biased expression, 6 were male-biased, and the remaining 15 genes either had no detectable expression or displayed unbiased expression. In the *S. purpurea* 15Z-SLR, 12 of the 33 single-copy genes were male-biased, while only three showed female-biased expression (for ten genes we did not obtain expression data; Fig. 4d; Table S23).

We calculated the TPM expression levels for 112 and 8 genes that are specific to the 15X- or 15Y-SLRs, respectively. The 15X-SLR specific genes exhibited a significant overall female-bias in expression, but one inserted (X insertion) gene (Saarb15aG0063700) showed male-biased expression (Table S24). We detected expression of only one 15Y-SLR-specific gene (Saarb15bG0048400) expressed, this suggests that 15Y-SLRs also accumulate male expressed genes. The genes shared only by the 15W and 15X-SLRs, or the 15Z and 15Y-SLRs, showed very distinct gene expression patterns, including (i) two 15X-specific genes that are not expressed, while the corresponding homologous genes of 15W expressed, (ii) one 15X-specific genes, and their 15W homologs, are female-biased or expressed only in females, (iii) neither 15X-specific gene (one) nor their 15W homolog is expressed, perhaps reflecting influences of repetitive sequences on expression, (iv) one 15X-specific gene is female-biased, while its 15W homologs is not expressed, (v) one 15X-specific is male-biased, and the 15W homolog is not expressed (Table S25). For the specific genes that shared by the 15Y and Z-SLR, these genes were found to be not expressed in 15Y-SLR but exhibited male-biased expression in 15Z. Overall, these results suggest that the original and/or accumulated genes on Z sex-linked region show masculinization, while the X-linked region ones show feminization.

## Discussion

### Expansion of the X sex-linked region in willows

In *S. arbutifolia*, the X-SLR (7.06 Mb) is considerably longer than the Y-linked region. The Y-SLR shows no evidence of degeneration, as only three pseudogenes were found. The large size of the X-SLR therefore probably reflects expansion of the X-SLR, consistent with the “extended region” that is evident in comparison with the *S. dunnii* chromosome 15, followed by changes such that its present repetitive sequence density is extremely high (86.50%) and the remainder is mostly duplicated genes. Although, as explained in the Introduction, X chromosomes may change in such ways (and some animal X chromosomes show intriguing properties that may reflect such changes), changes of the X are expected to be much more minor than those in Y-linked regions.

During sex chromosome evolution, repeats are expected to accumulate in non-recombining regions (Ming et al., 2011; Bachtrog, 2013; Charlesworth, 2017). These changes occur largely in the Y-linked regions, since the corresponding X-linked regions recombine in females, preventing most accumulation, though not all (see the Introduction). Therefore, the *S. arbutifolia* X situation remains unexplained. High densities of TEs and low gene content are expected in centromeric regions (Charlesworth et al., 1994), and these predictions are supported in willows (He et al., 2021). Our data for TE densities suggest that the SLRs may overlap with pericentromeric regions (Fig. S7). The “extended region” might therefore reflect such processes. An unknown type of repeat sequence shows a tendency to be inserted on X, and may preferentially insert into the X, similar to the preferences detected in *Silene latifolia* (Cermak et al., 2008; Filatov et al., 2009) and *Rumex acetosa* (Jesionek et al., 2021).

In addition to repetitive sequences, the 15X-SLR (but not the 15Y-SLR) includes a tandem array composed of 34 duplicate copies of a gene encoding a DExH-box ATP-dependent RNA helicase, which show female-biased expression in *S. arbutifolia*. DEAD-box ATP-dependent RNA helicase homologs have recently been identified in the candidate sex-determining region of grapevine (Badouin et al., 2020). Since the *S. purpurea* 15ZW-SLRs do not carry such a tandem array (Zhou et al., 2020), the duplications may have occurred in *S. arbutifolia* after the two species split, possibly because they have some female-specific function.

### Change in the heterogametic sex in willows

This study is the first to provide explicit details of a change in sex chromosome heterogamety in an angiosperm. We found two single-copy genes that may be completely sex-linked in the species with female heterogamety, and also in the species with male heterogamety, suggesting that the same gene or genes might be involved in sex-determination in both. The situation resembles the one studied in the platyfish, *Xiphophoris maculatus*, in which it has been proposed that the change might require only a change in copy number of a sex-determining gene, together with suitable dosage effects (Volff and Schartl, 2001). In an intriguing further similarity, the platyfish sex-determining region is also a complex and highly repetitive genome region (Tomaszkiewicz et al., 2014). Recombination in the region carrying the two shared genes with sex-specific alleles appears likely to have already been suppressed in the common ancestor of both species studied here, possibly due to proximity to the chromosome 15 centromere, which would also be consistent with the high repeat content we observe in these regions.

It is not surprising that the change in heterogamety involved the same ancestral chromosome, as this is predicted. This is because turnover is less likely when a species possesses highly degenerated Y or W chromosomes, which could lead to homozygotes of low fitness for these chromosomes (Bull, 1983). However, as suggested here, transitions involving a new sex-determining factor on the same chromosome do not result in such detrimental homozygotes. Consequently, even a slight advantage conferred by the new system could trigger turnover (Bull, 1983). For instance, the emergence of a more dominant female factor on the ancestral Y chromosome could favor the transition to a ZW system. It is also plausible that 15W derives from 15X, and 15Z from 15Y, as the X and W are feminizing chromosomes in both sex determination systems, whereas the Y and Z chromosome are expected to carry masculinizing factors. The model of (Volff and Schartl, 2001) suggests a way in which the same gene can act as a dominant masculinizing factor in an XY system and a recessive one in a ZW system, and it is intriguing that the systems studied here do indeed involve copy number differences. In these systems, it may become possible to test whether these, and suitable downstream dosage effects, are actually involved. 15X-SLR-specific genes showed predominantly female-biased expression, and a gene (Saarb15bG0048400, Table S24) inserted into the Y-SLR was also expressed male specifically, suggesting that the X- and Y-linked regions may respectively have accumulated feminizing and masculinizing factors. The single-copy genes with the topology II described in Fig. 3c generally show female-biased expression of the 15X alleles, and male-biased expression of 15Z ones (Fig. 4d, Table S23). Some single-copy sex-linked genes may therefore have evolved expression differences after the change in heterogamety.

## Methods

### Plant material

We collected young leaves from a male plant, B649AF, of *Salix arbutifolia* (15XY) for genome sequencing (Wang et al., 2023). Young leaf, catkin and stem samples for transcriptome sequencing were collected from B649AF in order to annotate genes. We made three biological replicates of the male and female catkins of B649AF and B649V, at full flowering for transcriptome sequencing to explore patterns of differential gene expression on sex chromosome of *S. arbutifolia*. Table S26 gives detailed information about the materials. Three species with published genome assemblies, *Populus trichocarpa* (19XY system) (Tuskan et al., 2006), *S. dunnii* (7XY system) (He et al., 2021) and *S. purpurea* (15ZW) (Zhou et al., 2020) (Fig. 1a) were included in some of our analyses.

### Genome sequencing

For Illumina PCR-free sequencing, total genomic DNA was extracted from B649AF using a Qiagen DNeasy Plant Mini kit following the manufacturer’s instructions (Qiagen).

PCR-free sequencing libraries were generated using the Illumina TruSeq DNA PCR-Free Library Preparation Kit (Illumina) following the manufacturer’s recommendations. For Hi-C and HiFi sequencing, total genomic DNA was extracted by the CTAB method (Chang et al., 1993). After quality assessment on an Agilent Bioanalyzer 2100 system, the PCR-free libraries were sequenced on an Illumina NovaSeq 6000platform by Beijing Novogene Bioinformatics Technology (hereafter Novogene).

The Hi-C library was prepared following a standard procedure (Wang et al., 2020). In brief, fresh leaves from B649AF were fixed in a 4% formaldehyde solution in MS buffer. Subsequently, crosslinked DNA was isolated from nuclei. The DpnII restriction enzyme was then used to digest the DNA, and the digested fragments were labelled with biotin, purified and ligated before sequencing. Hi-C libraries were controlled for quality and sequenced on an Illumina Hiseq X Ten platform by Novogene. The HiFi library for single-molecule real-time (SMRT) sequencing was constructed with an insert size of 15 kb using the SMRTbell Express Template Prep Kit 2.0. Briefly, the DNA was sheared using Diagenode Megaruptor system and concentrated with AMPure PB Beads (PacBio, CA, USA). The libraries were size selected on a BluePippin System. HiFi libraries were sequenced by Novogene on one SMRT cell using the PacBio Sequel II platform.

### RNA extraction and library preparation

Total RNA was extracted from young leaves, catkins and stems of B649AF and catkins of B649V using the Plant RNA Purification Reagent (Invitrogen) according to the manufacturer’s instructions. Genomic DNA was removed using DNase I (TaKara). RNA-seq transcriptome libraries were prepared using the TruSeq RNA sample preparation kit from Illumina, and sequencing was performed on an Illumina Novaseq 6000 by Novogene.

### Genome assembly

The CCS software (https://github.com/PacificBiosciences/ccs) was used to produce high-precision HiFi reads. The PacBio HiFi reads were used to assemble initial contigs using the Hifiasm pipeline v.0.16-r375 (Cheng et al., 2021) with default parameters, and to assemble haplotypes for subsequent analysis. We filtered the Hi-C reads using fastp (Chen et al., 2018), then mapped the clean reads to the assembled genome with Juicer (Durand et al., 2016), and finally assembled them using the 3d-DNA pipeline (Dudchenko et al., 2017). After a manual examination, we named the chromosome numbers of the assembly according to 19 chromosomes of *S. brachista* (Chen et al., 2019), based on the syntenic relationships of willows documented by (He et al., 2021).

The LR_Gapcloser (Xu et al., 2019b) was used to close gaps and improve the contiguity of genome assemblies based on HiFi reads. The telomere ends of most chromosomes are assembled, and have the telomere characteristic sequence (TTTAGGG)n, but some chromosomes are shorter or lack their telomeres. The assembly is therefore probably incomplete and the extension of the sequencing is insufficient, so the HiFi reads mapped to the chromosomes, and Hifiasm assembled reads from sequences near the telomere into separate contigs (Cheng et al., 2021). These contigs were therefore aligned with the chromosome sequences, to extend the chromosomes outward to complete their telomere sequences. We also employed NextPolish (Hu et al., 2020) to polish the assembly using Illumina short reads to improve base accuracy.

Redundans (Pryszcz and Gabaldon, 2016) was used to remove redundancy and contamination from external sources. This included: (i) aligning scattered contig sequences with chromosomes and organelle sequences to identify redundancy; (ii) identifying fragments or haplotigs with low average coverage depths in scattered contig sequences; (iii) identification of rDNA fragments scattered in the contig sequence. Finally, we manually checked possible redundant sequences and fragments, and removed them.

The GetOrganelle software (Jin et al., 2020) was used to assemble mitochondrial and chloroplast genomes based on Illumina short clean reads. The draft assembly was further evaluated by mapping the high-quality Illumina paired-end reads, HiFi reads, and RNA-Seq reads to the genome assembly using the BWA-MEM (v.0.7.8) algorithm (Li and Durbin, 2009), Minimap2 (Li, 2018), and HISAT2 (Kim et al., 2015). We also used BUSCO (v.4.1.2; http://busco.ezlab.org/) to check the completeness of the genome assembly.

### Annotation of repetitive sequences

A *de novo* TE library was generated using EDTA v1 (Ou et al., 2019) with parameter values “--sensitive 1 --anno 1”. Repeat elements were identified and classified using RepeatModeler (Flynn et al., 2020) to produce a repeat library. Then RepeatMasker (http://www.repeatmasker.org/RepeatMasker) was used to identify repeated regions in the genome, based on the library. The repeat-masked genome was subsequently used in gene annotation.

### Annotation of full-length LTR-RTs and estimation of insertion times

We annotated full-length LTR-RTs in our assembly and estimated their insertion times as described in (Xu et al., 2019a). Ltrharvest (Ellinghaus et al., 2008) and Ltrdigest (Steinbiss et al., 2009) were used to *de novo* predict full-length LTR-RTs in our assembly. LTR-RTs were then extracted and compared with *Gag-Pol* protein sequences in the REXdb database (Neumann et al., 2019). To estimate their insertion times, the LTRs of individual transposon insertions were aligned using MAFFT (Katoh and Standley, 2013), and divergence values between the 5’ and 3’-LTRs was estimated (SanMiguel et al., 1998; Ma and Bennetzen, 2004). The divergence values were corrected for saturation by Kimura’s 2-parameter method (Kimura, 1980), and insertion times were estimated from the values assuming a mutation rate of 2.5×10^−9^ substitutions year^−1^ per site, based on *Populus tremula* (Ingvarsson, 2008).

### Transcriptome assembly and gene annotation

The genome was annotated by combining evidence from transcriptome data, protein homology based, and *ab initio* prediction. We obtained three different transcriptomes by different methods: (i) RNA-seq reads *de novo* assembled with Trinity (Grabherr et al., 2011); (ii and iii) RNA-seq reads mapped to the reference genome using HISAT2 (Kim et al., 2015), and then assembled using either Trinity with genome-guided mode, or using StingTie (Pertea et al., 2015). CD-HIT (Fu et al., 2012) was used to remove short transcripts that were 95% covered by other transcripts with more than 95% identity in the three transcriptomes (redundant transcripts). The non-redundant transcripts were then used to compute assembly statistics for each method. PASA (Program to Assemble Spliced Alignment) (Haas et al., 2003) was used to obtain high-quality loci based on the transcriptome data. We randomly selected half of these loci as a training data set to train the AUGUSTUS (Stanke et al., 2008) gene modeller. The other half was used as the test data set, with five replicates for optimization. A total of 278,011 protein sequences were obtained from published data from 17 species of Salicaceae and *Arabidopsis thaliana* (Table S27), and used as reference proteins for homology-based gene annotation. MAKER2 pipeline (Cantarel et al., 2008) was used to annotate gene.

To annotate tRNA and rRNA sequences, we used tRNAScan-SE (Lowe and Eddy, 1997) and barrnap (https://github.com/tseemann/barrnap) (Lagesen et al., 2007), respectively, and discarded partial results. Other noncoding RNAs (ncRNAs) were identified by querying against the Rfam database (Nawrocki et al., 2015).

The functions of protein-coding genes were annotated based on three strategies. The annotated genes were aligned to proteins in the eggNOG databases (Powell et al., 2014) using eggNOG-Mapper (Huerta-Cepas et al., 2017), and assigned to GO (http://geneontology.org/) and KEGG (https://www.genome.jp/kegg/pathway.html) metabolic pathways to obtain functional information; (ii) the protein sequences were aligned with the Uniprot database (including Swiss-Prot and TrEMBL, https://www.uniprot.org/), NR (https://www.ncbi.nlm.nih.gov/), and *A. thaliana* reference genome sequence using Diamond (Buchfink et al., 2015) (with E value < 10^−5^ and identity > 30%); (iii) motifs and functional domains were identified by searching various domain libraries (ProDom, PRINTS, Pfam, SMART, PANTHER and PROSITE) using. Interproscan (Jones et al., 2014). Pseudopipe with default parameter settings was used to detect pseudogenes in the whole genome (Zhang and Yu, 2006). We also used blastp and blastn to identify pseudogenization of homologous genes within the X and Y-SLRs of *S. arbutifolia.* First, we performed blastp (-evalue 1e-5) to identify genes in 15X-SLR that did not have any peptide homologous genes in haplotype *a* (15X excluded). Next, we used blastn (-evalue 1e-5-word_size 8) to compare the coding sequences (CDS) of these genes with 15Y-SLR. Genes that showed alignment results with 15Y-SLR were classified as pseudogenes. In a second approach, in order to identify pseudogenes in 15Y-SLR, we extracted the possible pseudogenes identified by PseudoPipe in 15Y-SLR with the 15X gene IDs as the reference based. We used blastp (“-evalue 1e-5”) to blast the peptide sequences of candidate pseudogenes (with the 15X gene IDs) in 15Y-SLR against the peptide sequences of *S. arbutifolia* (15X excluded) and only kept pseudogenes that without hits on 15Y-SLR. Finally, we combined the pseudogenes that obtained by the two strategies but only kept one copy of duplicate results if there were. The same methods were applied to identify pseudogenes in 15X-SLR. The final SLR pseudogene datasets are the union of the two strategies.

### Identification of sex-associated SNPs and sex-linked region of *S. arbutifolia*

We also downloaded re-sequencing reads from 21 individual *S. arbutifolia* females and 20 males obtained by (Wang et al., 2023) to apply the chromosome quotient (CQ) method (Hall et al., 2013) to distinguish the 15X and 15Y haplotypes. CQ is the ratio of female to male alignments to a given reference sequence, using the stringent criterion that the entire read must align with zero mismatches. The CQ-calculate.pl software (https://sourceforge.net/projects/cqcalculate/files/CQ-calculate.pl/download) was used to map all female and male reads to two haplotypes (*a* and *b*), and calculate the CQ for each 50-kb nonoverlapping window, respectively. This can detect regions with different coverage in the DNA of the two sexes, such as degenerated Y-linked regions, in which X-linked genes are hemizygous in males, and also regions in which Y-linked sequences differ from their X-linked counterpart; this latter signal is less reliable, but offers an approach that can be used for young sex-linked regions in which degeneration is minor, or absent. In an XY system, CQ values should be close to two in X-SLR windows, but close to zero in Y-SLR windows.

The BWA-MEM algorithm of BWA 0.7.12 (Li and Durbin, 2009) was used to align the clean reads to the two assembled haplotypes (*a* and *b*). Samtools (Li et al., 2009) was used to extract primary alignments, sort, and merge the mapped data. Variants were jointly called for all individuals using Genome Analysis Toolkit (GATK) v. 4.1.8.1 (McKenna et al., 2010). GATK and VCFtools (Danecek et al., 2011) was used to select high-quality SNPs after hard filtering (QD< 2.0, FS > 60.0, MQ< 40.0, MQRankSum<−12.5, ReadPosRankSum <−8.0, SOR > 3.0). We (i) excluded sites with coverage exceedingly twice the mean depth at variant sites across all samples, (ii) retained only bi-allelic SNPs, and (iii) removed SNPs with missing information rate >10% and minor allele frequency <5%. A genome-wide association study (GWAS) was then performed using PLINK v1.07 (Purcell et al., 2007), and SNPs with associations at α < 0.05 after Bonferroni correction for multiple testing were identified as sex-associated. Weighted F_ST_ values between the sexes were calculated in 100-kb windows and 10-kb steps, using the Weir and Cockerham (Weir and Cockerham, 1984) estimator. A changepoint package (Killick and Eckley, 2014) was used to assess significance of differences in the mean and variance of the F_ST_ values between the sexes of sex chromosomes, using the function cpt.meanvar, algorithm PELT and penalty CROPS. Regions with significantly higher F_ST_ values than other parts of the chromosome are considered candidate SLRs (He et al., 2021).

High density of TEs and low gene content are expected in observed in the centromeric regions, therefore, we identified regions with such prominent patterns as putative centromeric regions.

### Micro-heteromorphism of X- and Y-SLRs of *S. arbutifolia*

We performed synteny analysis to compare the SLRs of the two sex chromosomes and their presumed ancestral autosome, chromosome 15 of outgroup *Salix* species (see below) using the Python version of MCScan (Tang et al., 2008) with “--cscore=.99”. To identify 15X-SLR-specific and 15Y-SLR-specific genes, we performed a reciprocal BLAST (Götz et al., 2008) of all peptide sequences in chromosome the 15X and 15Y regions with “blastp - evalue 1e-5”, and retained only gene pairs with identity ≥ 30% (Rost, 1999). 15X-SLR genes with no blast hits in 15Y-SLR were considered to be possibly 15X-SLR-specific, and vice versa for 15Y-SLR. Autosome 15 of two outgroup species, *S. dunnii* (He et al., 2021) and *P. trichocarpa* (Tuskan et al., 2006), were used to represent the likely state of the ancestral chromosome from which 15 X and Y evolved (Gulyaev et al., 2022). All the candidate 15X-SLR- and 15Y-SLR-specific genes were classified according to the following criteria: (i) 15X-specific (or 15Y-specific) genes with hits to chromosome 15 of *S. dunnii* and *P. trichocarpa* were categorized as “Y (or X) deletion”, (ii) genes that lacked hits to the chromosome 15 in both outgroup species were categorized as “X (or Y) insertion”. We employed blastp to identify duplicated genes within X-SLR and Y-SLR, based on their peptide sequences. We also used blastp to compare 15Y of *S. arbutifolia* and 15Z of *S. purpurea*.

### Tracing the heterogametic sex change in *Vetrix* clade

To trace the evolutionary history of 15XY and 15ZW in the *Vetrix* clade, we used the high-quality phased sex chromosome sequences of *S. arbutifolia* and *S. purpurea*. First, to test whether the sex-linked regions inherited similar sequences from their ancestors, we blasted all *S. arbutifolia* X-SLR and Y-SLR specific genes (see above) with the chromosome 15Z and 15W-SLRs of *S. purpurea* (Zhou et al., 2020), with “blastp - evalue 1e-5”. Second, we used OrthoFinder (Emms and Kelly, 2019) to identify the single-copy orthologs of 15X- and 15Y-linked genes of *S. arbutifolia*, 15Z- and 15W-linked genes of *S. purpurea* and chromosome 15 of *S. dunnii* (He et al., 2021; Wang et al., 2023). The MAGUS (Smirnov and Warnow, 2021) was used to align the sequences of each single-copy ortholog, before estimating gene trees using IQ-TREE (Nguyen et al., 2015), using *S. dunnii* as the outgroup to root the trees. ASTRAL version 5.7.8 (Zhang et al., 2018) was used to obtain species trees based on the topologies of the single-copy orthologs from the four SLRs using local posterior probabilities (Sayyari and Mirarab, 2016) to calculate clade support. To identify any ARR17-like gene duplicates in the SLRs, we used blastn to search the whole genomes of *S. arbutifolia* and *S. purpurea* with the Potri.019G133600 (Muller et al., 2020) sequence as the query, with the parameters “-evalue 1e-5-word_size 8”. We classified genes with all five CDS blastn results as intact ARR17-like gene duplicates, while blastn results lacking any of the five CDSs were considered partial duplicates (Wang et al., 2022). We extracted the CDS sequences of all intact and partial ARR17-like genes of *S. dunnii*, *S. purpurea*, and *S. arbutifolia*, aligned them using MAGUS, and then used IQ-TREE to estimate an ARR17-like gene tree, using orthologue in *A. thaliana* as outgroups to root the tree.

### Identification of possible evolutionary strata in X and Z chromosomes

We distinguished between different evolutionary strata according to the topologies of the single-copy orthologs of X-, Y-, Z-, and W-SLR genes of *S. arbutifolia* and *S. purpurea*. These topologies reflect the times when X-Y or W-Z recombination ceased (Handley et al., 2004). The early and late recombination suppressed genes along the sex chromosome will exhibit different topologies and thus can be used to infer evolutionary strata (Handley et al., 2004; Zhang et al., 2022a). Using the yn00 function in PAML (Yang, 2007), we calculated synonymous site divergence (Ks) values based on orthologous pairs of 15X- and 15Y-SLRs (or 15W- and 15Z-SLRs) identified by reciprocal blast (Götz et al., 2008) with“blastp - evalue 1e- 5 - max_target_seqs 1” and identify > 70 % (Sievers et al., 2011). Lower Ks values indicate younger strata.

### Gene expression

After assessing the RNA-seq data quality with FASTQC v0.12.0 (https://www.bioinformatics.babraham.ac.uk/projects/fastqc/), we used FASTP (Chen et al., 2018) to remove adaptor sequences and trim the raw reads from male and female samples (Table S26). Then we mapped the clean RNA-seq reads to haplotype *a* (15X kept) and 15Y in the assembled genome (identified as explained above) using HISAT2 (Kim et al., 2015) and assigned counts using featureCounts (Liao et al., 2014). Then, after filtering out unexpressed genes (counts=0 in all samples, excluding non-mRNA), we converted the read counts to TPM (transcripts per million reads). Differential expression analyses between male and female groups (with three biological replicates per group) were performed using the DESeq2 R package (1.16.1) (Love et al., 2014). Genes with corrected *P*-values < 0.05 and |log2FoldChange| >1 (absolute value of log2FoldChange > 1) as identified by DESeq2 were classified as differentially expressed (DEGs). We also included gene expression data of male and female catkins from *S. purpurea* (Hyden et al., 2021) to help interpret the differential gene expression patterns.

## Acknowledgements

This study was financially supported by the National Natural Science Foundation of China (grant no. 32171813) and Special Fund for Scientific Research of Shanghai Landscaping & City Appearance Administrative Bureau (grant no. G232403). We are grateful to Yuan Wang, Zhi-Qing Xue, and Xin-jie Cai for their suggestions on our analysis, and Xiao-Bo An for sampling.

## Competing interests

The authors declare no competing interests.

## Author contributions

Yi Wang and Li He planned and designed the research; Yi Wang, Li He, Rengang Zhang and Guangnan Gong analyzed the data; Yi Wang, Li He, Elvira Hörandl, Zhixiang Zhang, and Deborah Charlesworth wrote the paper.

## Data availability

Sequencing data for genome assembly and annotation can be downloaded from the National Center for Biotechnology Information (NCBI) under the BioProject accession number PRJNA882493. The genome assembly and annotation can be downloaded from the National Genomics Data Center (NGDC) under the BioProject accession number PRJCA016000. Transcriptome data of male and female catkins of B649AF and B649V can be downloaded from the NCBI Sequence Read Archive (SRA) under the BioProject accession number PRJNA990983.

## Supporting information

Table S1: Sequencing statistics of WGS-HiFi, WGS-Illumina paired-end sequences, Hi-C and RNASeq datasets.

Table S2: Sequencing data sets map to genome ratio and coverage percentage statistics.

Table S3: Summary of repeat contents of the genome of *Salix arbutifolia*.

Table S4: The statistics for full-length long terminal repeat-retrotransposons (LTR-RTs) of *Salix arbutifolia* genome.

Table S5: Source statistics for gene annotation.

Table S6: Statistics of RNAs of the genome of *Salix arbutifolia*.

Table S7: Functional annotation of the predicted genes of *Salix arbutifolia*.

Table S8: Summary of mapping results of 41 samples of *Salix arbutifolia* using haplotype *a* as reference.

Table S9: Summary of mapping results of 41 samples of *Salix arbutifolia* using haplotype *b* as reference.

Table S10: Statistics of significantly sex associated SNPs and sex-linked genes in the *Salix arbutifolia* 15X-SLR.

Table S11: Pseudogenes of sex chromosome 15X and Y of *S. arbutifolia*.

Table S12: The type and size of the repeat sequences in the 15X-SLR extended region.

Table S13: Statistics of gene duplicates ID and length of extended region of 15X-SLRs.

Table S14: Thirty-four tandem genes in extended region of 15X-SLR.

Table S15: 15X-specific genes relative to 15Y-SLR obtained by reciprocal BLAST.

Table S16: 15Y-specific genes relative to 15X-SLR obtained by reciprocal BLAST.

Table S17: Ka, Ks values between homologous gene pairs of chromosomes 15X and 15Y of *S. arbutifolia*. Different color areas represent different evolutionary stratums.

Table S18: Ks value of different evolutionary stratums of *S. purpurea* 15 Z and W-SLRs.

Table S19: Length difference between *S. arbutifolia* 15Y and *S. purpurea* 15Z composition.

Table S20: Transcriptome data quality control and mapping results.

Table S21: Statistics on the number of sex-biased genes on 19 chromosomes of *S. arbutifolia*.

Table S22: The gene expression of a tandem array (34 genes) within the 15X-SLR.

Table S23: The gene expression patterns for 33 single-copy genes representing topologies I and II.

Table S24: The gene expression patterns of 15X- and 15Y-specific genes.

Table S25: The specific genes expression patterns that shared only by 15W and 15X-SLR or 15Z and 15Y-SLR.

Table S26: Details of plant materials used in this study.

Table S27: Details of 17 species of Salicaceae and *Arabidopsis thaliana* used as references for homology-based gene annotation.

Figure S1: All the CQ results of haplotype *a* of *S. arbutifolia*.

Figure S2: All the CQ results of haplotype *b* of *S. arbutifolia*.

Figure S3: All the F_ST_ results of haplotype *a* of *S. arbutifolia*.

Figure S4: All the F_ST_ results of haplotype *b* of *S. arbutifolia*.

Figure S5: Quantile-Quantile (Q-Q) plots of observed and expected GWAS *P*-values of *Salix arbutifolia* haplotype *a* and results of GWAS between SNPs and the sexes of 39 individuals use *Salix arbutifolia* haplotype *a* as reference.

Figure S6: Quantile-Quantile (Q-Q) plots of observed and expected GWAS *P*-values of *Salix arbutifolia* haplotype *b* and results of GWAS between SNPs and the sexes of 39 individuals use *Salix arbutifolia* haplotype *b* as reference.

Figure S7: Gene density, transposable element (TE) density, LTR-Copia density and LTR-Gypsy density of genome *Salix arbutifolia*.

Figure S8: The gene tree of all intact and partial ARR17-like genes of *S. dunnii*, *S. purpurea*, and *S. arbutifolia*.

Figure S9: Distribution of Ks values of *S. arbutifolia* 15X-15Y gene pairs and *S. purpurea* 15Z-SLR-15W-SLR gene pairs.

Figure S10: Collinearity analysis using *S. dunnii* autosome 15 and *S. purpurea* 15Z and W chromosomes.

Figure S11: Collinearity analysis using *S. purpurea* 15Z*, S. arbutifolia* 15Y *and S. dunnii* autosome 15.

## Notes

### Competing Interest Statement

The authors have declared no competing interest.

## References

Almeida, P., Proux-Wera, E., Churcher, A., Soler, L., Dainat, J., Pucholt, P., Nordlund, J., Martin, T., Ronnberg-Wastljung, A.C., Nystedt, B., Berlin, S., and Mank, J.E. (2020). Genome assembly of the basket willow, *Salix viminalis*, reveals earliest stages of sex chromosome expansion. BMC Biol 18, 78.

Bachtrog, D. (2013). Y-chromosome evolution: emerging insights into processes of Y-chromosome degeneration. Nat Rev Genet 14, 113–124.

Badouin, H., Velt, A., Gindraud, F., Flutre, T., Dumas, V., Vautrin, S., Marande, W., Corbi, J., Sallet, E., Ganofsky, J., Santoni, S., Guyot, D., Ricciardelli, E., Jepsen, K., Käfer, J., Berges, H., Duchêne, E., Picard, F., Hugueney, P., Tavares, R., Bacilieri, R., Rustenholz, C., and Marais, G.A.B. (2020). The wild grape genome sequence provides insights into the transition from dioecy to hermaphroditism during grape domestication. Genome Biol 21, 223.

Bergero, R., and Charlesworth, D. (2009). The evolution of restricted recombination in sex chromosomes. Trends Ecol Evol 24, 94–102.

Bergero, R., and Charlesworth, D. (2011). Preservation of the Y transcriptome in a 10-million-year-old plant sex chromosome system. Curr Biol 21, 1470–1474.

Bergero, R., Forrest, A., and Charlesworth, D. (2008). Active miniature transposons from a plant genome and its nonrecombining Y chromosome. Genetics 178, 1085–1092.

Buchfink, B., Xie, C., and Huson, D.H. (2015). Fast and sensitive protein alignment using DIAMOND. Nat Methods 12, 59–60.

Bull, J.J. (1983). Evolution of sex determining mechanisms. (The Benjamin/Cummings Publishing Company, Inc.).

Bull, J.J. (1985). Sex determining mechanisms: an evolutionary perspective. Experientia 41, 1285–1296.

Cantarel, B.L., Korf, I., Robb, S.M., Parra, G., Ross, E., Moore, B., Holt, C., Sánchez Alvarado, A., and Yandell, M. (2008). MAKER: an easy-to-use annotation pipeline designed for emerging model organism genomes. Genome Res 18, 188–196.

Cermak, T., Kubat, Z., Hobza, R., Koblizkova, A., Widmer, A., Macas, J., Vyskot, B., and Kejnovsky, E. (2008). Survey of repetitive sequences in *Silene latifolia* with respect to their distribution on sex chromosomes. Chromosome Res 16, 961–976.

Chang, S., Puryear, J., and Cairney, J. (1993). A simple and efficient method for isolating RNA from pine trees. Plant Molecular Biology Reporter 11, 113–116.

Charlesworth, B., and Charlesworth, D. (1978). A Model for the Evolution of Dioecy and Gynodioecy. The American Naturalist 112, 975–997.

Charlesworth, B., Sniegowski, P., and Stephan, W. (1994). The evolutionary dynamics of repetitive DNA in eukaryotes. Nature 371, 215–220.

Charlesworth, D. (2017). Evolution of recombination rates between sex chromosomes. Philos Trans R Soc Lond B Biol Sci 372.

Charlesworth, D., Charlesworth, B., and Marais, G. (2005). Steps in the evolution of heteromorphic sex chromosomes. Heredity (Edinb) 95, 118–128.

Chen, J.H., Huang, Y., Brachi, B., Yun, Q.Z., Zhang, W., Lu, W., Li, H.N., Li, W.Q., Sun, X.D., Wang, G.Y., He, J., Zhou, Z., Chen, K.Y., Ji, Y.H., Shi, M.M., Sun, W.G., Yang, Y.P., Zhang, R.G., Abbott, R.J., and Sun, H. (2019). Genome-wide analysis of Cushion willow provides insights into alpine plant divergence in a biodiversity hotspot. Nat Commun 10, 5230.

Chen, S., Zhou, Y., Chen, Y., and Gu, J. (2018). fastp: an ultra-fast all-in-one FASTQ preprocessor. Bioinformatics 34, i884–i890.

Cheng, H., Concepcion, G.T., Feng, X., Zhang, H., and Li, H. (2021). Haplotype-resolved de novo assembly using phased assembly graphs with hifiasm. Nat Methods 18, 170–175.

Chibalina, M.V., and Filatov, D.A. (2011). Plant Y chromosome degeneration is retarded by haploid purifying selection. Curr Biol 21, 1475–1479.

Crowson, D., Barrett, S.C.H., and Wright, S.I. (2017). Purifying and Positive Selection Influence Patterns of Gene Loss and Gene Expression in the Evolution of a Plant Sex Chromosome System. Mol Biol Evol 34, 1140–1154.

Danecek, P., Auton, A., Abecasis, G., Albers, C.A., Banks, E., DePristo, M.A., Handsaker, R.E., Lunter, G., Marth, G.T., Sherry, S.T., McVean, G., Durbin, R., and Genomes Project Analysis, G. (2011). The variant call format and VCFtools. Bioinformatics 27, 2156–2158.

Dudchenko, O., Batra, S.S., Omer, A.D., Nyquist, S.K., Hoeger, M., Durand, N.C., Shamim, M.S., Machol, I., Lander, E.S., Aiden, A.P., and Aiden, E.L. (2017). De novo assembly of the *Aedes aegypti* genome using Hi-C yields chromosome-length scaffolds. Science 356, 92–95.

Durand, N.C., Shamim, M.S., Machol, I., Rao, S.S., Huntley, M.H., Lander, E.S., and Aiden, E.L. (2016). Juicer Provides a One-Click System for Analyzing Loop-Resolution Hi-C Experiments. Cell Syst 3, 95–98.

Ellinghaus, D., Kurtz, S., and Willhoeft, U. (2008). LTRharvest, an efficient and flexible software for de novo detection of LTR retrotransposons. BMC Bioinformatics 9, 18.

Emms, D.M., and Kelly, S. (2019). OrthoFinder: phylogenetic orthology inference for comparative genomics. Genome Biol 20, 238.

Filatov, D.A., Howell, E.C., Groutides, C., and Armstrong, S.J. (2009). Recent spread of a retrotransposon in the *Silene latifolia* genome, apart from the Y chromosome. Genetics 181, 811–817.

Flynn, J.M., Hubley, R., Goubert, C., Rosen, J., Clark, A.G., Feschotte, C., and Smit, A.F. (2020). RepeatModeler2 for automated genomic discovery of transposable element families. Proc Natl Acad Sci U S A 117, 9451–9457.

Fraisse, C., Picard, M.A.L., and Vicoso, B. (2017). The deep conservation of the Lepidoptera Z chromosome suggests a non-canonical origin of the W. Nat Commun 8, 1486.

Fu, L., Niu, B., Zhu, Z., Wu, S., and Li, W. (2012). CD-HIT: accelerated for clustering the next-generation sequencing data. Bioinformatics 28, 3150–3152.

Furman, B.L.S., Cauret, C.M.S., Knytl, M., Song, X.Y., Premachandra, T., Ofori-Boateng, C., Jordan, D.C., Horb, M.E., and Evans, B.J. (2020). A frog with three sex chromosomes that co-mingle together in nature: *Xenopus tropicalis* has a degenerate W and a Y that evolved from a Z chromosome. PLoS Genet 16, e1009121.

Götz, S., García-Gómez, J.M., Terol, J., Williams, T.D., Nagaraj, S.H., Nueda, M.J., Robles, M., Talón, M., Dopazo, J., and Conesa, A. (2008). High-throughput functional annotation and data mining with the Blast2GO suite. Nucleic Acids Res 36, 3420–3435.

Grabherr, M.G., Haas, B.J., Yassour, M., Levin, J.Z., Thompson, D.A., Amit, I., Adiconis, X., Fan, L., Raychowdhury, R., Zeng, Q., Chen, Z., Mauceli, E., Hacohen, N., Gnirke, A., Rhind, N., di Palma, F., Birren, B.W., Nusbaum, C., Lindblad-Toh, K., Friedman, N., and Regev, A. (2011). Full-length transcriptome assembly from RNA-Seq data without a reference genome. Nat Biotechnol 29, 644–652.

Gulyaev, S., Cai, X.J., Guo, F.Y., Kikuchi, S., Applequist, W.L., Zhang, Z.X., Horandl, E., and He, L. (2022). The phylogeny of *Salix* revealed by whole genome re-sequencing suggests different sex-determination systems in major groups of the genus. Ann Bot 129, 485–498.

Haas, B.J., Delcher, A.L., Mount, S.M., Wortman, J.R., Smith, R.K., Jr., Hannick, L.I., Maiti, R., Ronning, C.M., Rusch, D.B., Town, C.D., Salzberg, S.L., and White, O. (2003). Improving the *Arabidopsis* genome annotation using maximal transcript alignment assemblies. Nucleic Acids Res 31, 5654–5666.

Hall, A.B., Qi, Y., Timoshevskiy, V., Sharakhova, M.V., Sharakhov, I.V., and Tu, Z. (2013). Six novel Y chromosome genes in *Anopheles* mosquitoes discovered by independently sequencing males and females. BMC Genomics 14, 273.

Handley, L.J., Ceplitis, H., and Ellegren, H. (2004). Evolutionary strata on the chicken Z chromosome: implications for sex chromosome evolution. Genetics 167, 367–376.

He, L., Jia, K.H., Zhang, R.G., Wang, Y., Shi, T.L., Li, Z.C., Zeng, S.W., Cai, X.J., Wagner, N.D., Horandl, E., Muyle, A., Yang, K., Charlesworth, D., and Mao, J.F. (2021). Chromosome-scale assembly of the genome of *Salix dunnii* reveals a male-heterogametic sex determination system on chromosome 7. Mol Ecol Resour 21, 1966–1982.

Hu, J., Fan, J., Sun, Z., and Liu, S. (2020). NextPolish: a fast and efficient genome polishing tool for long-read assembly. Bioinformatics 36, 2253–2255.

Hu, N., Sanderson, B., Guo, M., Feng, G., Gambhir, D., Hale, H., Wang, D., Hyden, B., Liu, J., Ma, T., DiFazio, S., Smart, L., and Olson, M. (2022). An unusual origin of a ZW sex chromosome system (Research Square).

Huerta-Cepas, J., Forslund, K., Coelho, L.P., Szklarczyk, D., Jensen, L.J., von Mering, C., and Bork, P. (2017). Fast Genome-Wide Functional Annotation through Orthology Assignment by eggNOG-Mapper. Mol Biol Evol 34, 2115–2122.

Hyden, B., Zou, J., Wilkerson, D.G., Carlson, C.H., Robles, A.R., DiFazio, S.P., and Smart, L.B. (2023). Structural variation of a sex-linked region confers monoecy and implicates GATA15 as a master regulator of sex in *Salix purpurea*. New Phytol.

Hyden, B., Carlson, C.H., Gouker, F.E., Schmutz, J., Barry, K., Lipzen, A., Sharma, A., Sandor, L., Tuskan, G.A., Feng, G., Olson, M.S., DiFazio, S.P., and Smart, L.B. (2021). Integrative genomics reveals paths to sex dimorphism in *Salix purpurea* L. Hortic Res 8, 170.

Ieda, R., Hosoya, S., Tajima, S., Atsumi, K., Kamiya, T., Nozawa, A., Aoki, Y., Tasumi, S., Koyama, T., Nakamura, O., Suzuki, Y., and Kikuchi, K. (2018). Identification of the sex-determining locus in grass puffer (*Takifugu niphobles*) provides evidence for sex-chromosome turnover in a subset of Takifugu species. PLoS One 13, e0190635.

Ingvarsson, P.K. (2008). Multilocus patterns of nucleotide polymorphism and the demographic history of *Populus tremula*. Genetics 180, 329–340.

Jesionek, W., Bodlakova, M., Kubat, Z., Cegan, R., Vyskot, B., Vrana, J., Safar, J., Puterova, J., and Hobza, R. (2021). Fundamentally different repetitive element composition of sex chromosomes in *Rumex acetosa*. Ann Bot 127, 33–47.

Jin, J.J., Yu, W.B., Yang, J.B., Song, Y., dePamphilis, C.W., Yi, T.S., and Li, D.Z. (2020). GetOrganelle: a fast and versatile toolkit for accurate de novo assembly of organelle genomes. Genome Biol 21, 241.

Jones, P., Binns, D., Chang, H.Y., Fraser, M., Li, W., McAnulla, C., McWilliam, H., Maslen, J., Mitchell, A., Nuka, G., Pesseat, S., Quinn, A.F., Sangrador-Vegas, A., Scheremetjew, M., Yong, S.Y., Lopez, R., and Hunter, S. (2014). InterProScan 5: genome-scale protein function classification. Bioinformatics 30, 1236–1240.

Kabir, A., Ieda, R., Hosoya, S., Fujikawa, D., Atsumi, K., Tajima, S., Nozawa, A., Koyama, T., Hirase, S., Nakamura, O., Kadota, M., Nishimura, O., Kuraku, S., Nakamura, Y., Kobayashi, H., Toyoda, A., Tasumi, S., and Kikuchi, K. (2022). Repeated translocation of a supergene underlying rapid sex chromosome turnover in Takifugu pufferfish. Proc Natl Acad Sci U S A 119, e2121469119.

Katoh, K., and Standley, D.M. (2013). MAFFT multiple sequence alignment software version 7: improvements in performance and usability. Mol Biol Evol 30, 772–780.

Killick, R., and Eckley, I.A. (2014). changepoint: An R Package for Changepoint Analysis. Journal of Statistical Software 58, 1–19.

Kim, D., Langmead, B., and Salzberg, S.L. (2015). HISAT: a fast spliced aligner with low memory requirements. Nat Methods 12, 357–360.

Kimura, M. (1980). A simple method for estimating evolutionary rates of base substitutions through comparative studies of nucleotide sequences. J Mol Evol 16, 111–120.

Lagesen, K., Hallin, P., Rødland, E.A., Staerfeldt, H.H., Rognes, T., and Ussery, D.W. (2007). RNAmmer: consistent and rapid annotation of ribosomal RNA genes. Nucleic Acids Res 35, 3100–3108.

Lahn, B.T., and Page, D.C. (1999). Four evolutionary strata on the human X chromosome. Science 286, 964–967.

Li, H. (2018). Minimap2: pairwise alignment for nucleotide sequences. Bioinformatics 34, 3094–3100.

Li, H., and Durbin, R. (2009). Fast and accurate short read alignment with Burrows-Wheeler transform. Bioinformatics 25, 1754–1760.

Li, H., Handsaker, B., Wysoker, A., Fennell, T., Ruan, J., Homer, N., Marth, G., Abecasis, G., and Durbin, R. (2009). The Sequence Alignment/Map format and SAMtools. Bioinformatics 25, 2078–2079.

Li, Y., Wang, D., Wang, W., Yang, W., Gao, J., Zhang, W., Shan, L., Kang, M., Chen, Y., and Ma, T. (2022). A chromosome-level *Populus qiongdaoensis* genome assembly provides insights into tropical adaptation and a cryptic turnover of sex determination. Mol Ecol.

Liao, Y., Smyth, G.K., and Shi, W. (2014). featureCounts: an efficient general purpose program for assigning sequence reads to genomic features. Bioinformatics 30, 923–930.

Love, M.I., Huber, W., and Anders, S. (2014). Moderated estimation of fold change and dispersion for RNA-seq data with DESeq2. Genome Biol 15, 550.

Lowe, T.M., and Eddy, S.R. (1997). tRNAscan-SE: a program for improved detection of transfer RNA genes in genomic sequence. Nucleic Acids Res 25, 955–964.

Ma, J., and Bennetzen, J.L. (2004). Rapid recent growth and divergence of rice nuclear genomes. Proc Natl Acad Sci U S A 101, 12404–12410.

Ma, W.J., Veltsos, P., Sermier, R., Parker, D.J., and Perrin, N. (2018). Evolutionary and developmental dynamics of sex-biased gene expression in common frogs with proto-Y chromosomes. Genome Biol 19, 156.

Mahrez, W., Shin, J., Muñoz-Viana, R., Figueiredo, D.D., Trejo-Arellano, M.S., Exner, V., Siretskiy, A., Gruissem, W., Köhler, C., and Hennig, L. (2016). BRR2a Affects Flowering Time via FLC Splicing. PLOS Genetics 12, e1005924.

McKenna, A., Hanna, M., Banks, E., Sivachenko, A., Cibulskis, K., Kernytsky, A., Garimella, K., Altshuler, D., Gabriel, S., Daly, M., and DePristo, M.A. (2010). The Genome Analysis Toolkit: a MapReduce framework for analyzing next-generation DNA sequencing data. Genome Res 20, 1297–1303.

Mensah, M.A., Hestand, M.S., Larmuseau, M.H., Isrie, M., Vanderheyden, N., Declercq, M., Souche, E.L., Van Houdt, J., Stoeva, R., Van Esch, H., Devriendt, K., Voet, T., Decorte, R., Robinson, P.N., and Vermeesch, J.R. (2014). Pseudoautosomal region 1 length polymorphism in the human population. PLoS Genet 10, e1004578.

Ming, R., Bendahmane, A., and Renner, S.S. (2011). Sex chromosomes in land plants. Annu Rev Plant Biol 62, 485–514.

Miura, I. (2007). An evolutionary witness: the frog *rana rugosa* underwent change of heterogametic sex from XY male to ZW female. Sex Dev 1, 323–331.

Mrnjavac, A., Khudiakova, K.A., Barton, N.H., and Vicoso, B. (2023). Slower-X: reduced efficiency of selection in the early stages of X chromosome evolution. Evol Lett 7, 4–12.

Muller, N.A., Kersten, B., Leite Montalvao, A.P., Mahler, N., Bernhardsson, C., Brautigam, K., Carracedo Lorenzo, Z., Hoenicka, H., Kumar, V., Mader, M., Pakull, B., Robinson, K.M., Sabatti, M., Vettori, C., Ingvarsson, P.K., Cronk, Q., Street, N.R., and Fladung, M. (2020). A single gene underlies the dynamic evolution of poplar sex determination. Nat Plants 6, 630–637.

Nawrocki, E.P., Burge, S.W., Bateman, A., Daub, J., Eberhardt, R.Y., Eddy, S.R., Floden, E.W., Gardner, P.P., Jones, T.A., Tate, J., and Finn, R.D. (2015). Rfam 12.0: updates to the RNA families database. Nucleic Acids Res 43, D130–137.

Neumann, P., Novák, P., Hoštáková, N., and Macas, J. (2019). Systematic survey of plant LTR-retrotransposons elucidates phylogenetic relationships of their polyprotein domains and provides a reference for element classification. Mob DNA 10, 1.

Nguyen, L.T., Schmidt, H.A., von Haeseler, A., and Minh, B.Q. (2015). IQ-TREE: a fast and effective stochastic algorithm for estimating maximum-likelihood phylogenies. Mol Biol Evol 32, 268–274.

Ogata, M., Lambert, M., Ezaz, T., and Miura, I. (2018). Reconstruction of female heterogamety from admixture of XX-XY and ZZ-ZW sex-chromosome systems within a frog species. Mol Ecol 27, 4078–4089.

Ou, S., Su, W., Liao, Y., Chougule, K., Agda, J.R.A., Hellinga, A.J., Lugo, C.S.B., Elliott, T.A., Ware, D., Peterson, T., Jiang, N., Hirsch, C.N., and Hufford, M.B. (2019). Benchmarking transposable element annotation methods for creation of a streamlined, comprehensive pipeline. Genome Biol 20, 275.

Papadopulos, A.S., Chester, M., Ridout, K., and Filatov, D.A. (2015). Rapid Y degeneration and dosage compensation in plant sex chromosomes. Proc Natl Acad Sci U S A 112, 13021–13026.

Pertea, M., Pertea, G.M., Antonescu, C.M., Chang, T.C., Mendell, J.T., and Salzberg, S.L. (2015). StringTie enables improved reconstruction of a transcriptome from RNA-seq reads. Nat Biotechnol 33, 290–295.

Picard, M.A.L., Cosseau, C., Ferre, S., Quack, T., Grevelding, C.G., Coute, Y., and Vicoso, B. (2018). Evolution of gene dosage on the Z-chromosome of schistosome parasites. Elife 7.

Powell, S., Forslund, K., Szklarczyk, D., Trachana, K., Roth, A., Huerta-Cepas, J., Gabaldón, T., Rattei, T., Creevey, C., Kuhn, M., Jensen, L.J., von Mering, C., and Bork, P. (2014). eggNOG v4.0: nested orthology inference across 3686 organisms. Nucleic Acids Res 42, D231–239.

Prentout, D., Stajner, N., Cerenak, A., Tricou, T., Brochier-Armanet, C., Jakse, J., Kafer, J., and Marais, G.A.B. (2021). Plant genera *Cannabis* and *Humulus* share the same pair of well-differentiated sex chromosomes. New Phytol 231, 1599–1611.

Pryszcz, L.P., and Gabaldon, T. (2016). Redundans: an assembly pipeline for highly heterozygous genomes. Nucleic Acids Res 44, e113.

Purcell, S., Neale, B., Todd-Brown, K., Thomas, L., Ferreira, M.A., Bender, D., Maller, J., Sklar, P., de Bakker, P.I., Daly, M.J., and Sham, P.C. (2007). PLINK: a tool set for whole-genome association and population-based linkage analyses. Am J Hum Genet 81, 559–575.

Rost, B. (1999). Twilight zone of protein sequence alignments. Protein Engineering, Design and Selection 12, 85–94.

Rozen, S., Skaletsky, H., Marszalek, J.D., Minx, P.J., Cordum, H.S., Waterston, R.H., Wilson, R.K., and Page, D.C. (2003). Abundant gene conversion between arms of palindromes in human and ape Y chromosomes. Nature 423, 873–876.

SanMiguel, P., Gaut, B.S., Tikhonov, A., Nakajima, Y., and Bennetzen, J.L. (1998). The paleontology of intergene retrotransposons of maize. Nat Genet 20, 43–45.

Saunders, P.A., Neuenschwander, S., and Perrin, N. (2018). Sex chromosome turnovers and genetic drift: a simulation study. J Evol Biol 31, 1413–1419.

Sayyari, E., and Mirarab, S. (2016). Fast Coalescent-Based Computation of Local Branch Support from Quartet Frequencies. Mol Biol Evol 33, 1654–1668.

Sievers, F., Wilm, A., Dineen, D., Gibson, T.J., Karplus, K., Li, W., Lopez, R., McWilliam, H., Remmert, M., Söding, J., Thompson, J.D., and Higgins, D.G. (2011). Fast, scalable generation of high-quality protein multiple sequence alignments using Clustal Omega. Mol Syst Biol 7, 539.

Smirnov, V., and Warnow, T. (2021). MAGUS: Multiple sequence Alignment using Graph clUStering. Bioinformatics 37, 1666–1672.

Stanke, M., Diekhans, M., Baertsch, R., and Haussler, D. (2008). Using native and syntenically mapped cDNA alignments to improve de novo gene finding. Bioinformatics 24, 637–644.

Steinbiss, S., Willhoeft, U., Gremme, G., and Kurtz, S. (2009). Fine-grained annotation and classification of de novo predicted LTR retrotransposons. Nucleic Acids Res 37, 7002–7013.

Tang, H., Bowers, J.E., Wang, X., Ming, R., Alam, M., and Paterson, A.H. (2008). Synteny and collinearity in plant genomes. Science 320, 486–488.

Tomaszkiewicz, M., Chalopin, D., Schartl, M., Galiana, D., and Volff, J.N. (2014). A multicopy Y-chromosomal SGNH hydrolase gene expressed in the testis of the platyfish has been captured and mobilized by a Helitron transposon. BMC Genet 15, 44.

Tuskan, G.A., Difazio, S., Jansson, S., Bohlmann, J., Grigoriev, I., Hellsten, U., Putnam, N., Ralph, S., Rombauts, S., Salamov, A., Schein, J., Sterck, L., Aerts, A., Bhalerao, R.R., Bhalerao, R.P., Blaudez, D., Boerjan, W., Brun, A., Brunner, A., Busov, V., Campbell, M., Carlson, J., Chalot, M., Chapman, J., Chen, G.L., Cooper, D., Coutinho, P.M., Couturier, J., Covert, S., Cronk, Q., Cunningham, R., Davis, J., Degroeve, S., Déjardin, A., Depamphilis, C., Detter, J., Dirks, B., Dubchak, I., Duplessis, S., Ehlting, J., Ellis, B., Gendler, K., Goodstein, D., Gribskov, M., Grimwood, J., Groover, A., Gunter, L., Hamberger, B., Heinze, B., Helariutta, Y., Henrissat, B., Holligan, D., Holt, R., Huang, W., Islam-Faridi, N., Jones, S., Jones-Rhoades, M., Jorgensen, R., Joshi, C., Kangasjärvi, J., Karlsson, J., Kelleher, C., Kirkpatrick, R., Kirst, M., Kohler, A., Kalluri, U., Larimer, F., Leebens-Mack, J., Leplé, J.C., Locascio, P., Lou, Y., Lucas, S., Martin, F., Montanini, B., Napoli, C., Nelson, D.R., Nelson, C., Nieminen, K., Nilsson, O., Pereda, V., Peter, G., Philippe, R., Pilate, G., Poliakov, A., Razumovskaya, J., Richardson, P., Rinaldi, C., Ritland, K., Rouzé, P., Ryaboy, D., Schmutz, J., Schrader, J., Segerman, B., Shin, H., Siddiqui, A., Sterky, F., Terry, A., Tsai, C.J., Uberbacher, E., Unneberg, P., Vahala, J., Wall, K., Wessler, S., Yang, G., Yin, T., Douglas, C., Marra, M., Sandberg, G., Van de Peer, Y., and Rokhsar, D. (2006). The genome of black cottonwood, *Populus trichocarpa* (Torr. & Gray). Science 313, 1596–1604.

van Doorn, G.S., and Kirkpatrick, M. (2010). Transitions between male and female heterogamety caused by sex-antagonistic selection. Genetics 186, 629–645.

Volff, J.N., and Schartl, M. (2001). Variability of genetic sex determination in poeciliid fishes. Genetica 111, 101–110.

Wang, D., Li, Y., Li, M., Yang, W., Ma, X., Zhang, L., Wang, Y., Feng, Y., Zhang, Y., Zhou, R., Sanderson, B.J., Keefover-Ring, K., Yin, T., Smart, L.B., DiFazio, S.P., Liu, J., Olson, M., and Ma, T. (2022). Repeated turnovers keep sex chromosomes young in willows. Genome Biol 23, 200.

Wang, H., Sun, S., Ge, W., Zhao, L., Hou, B., Wang, K., Lyu, Z., Chen, L., Xu, S., Guo, J., Li, M., Su, P., Li, X., Wang, G., Bo, C., Fang, X., Zhuang, W., Cheng, X., Wu, J., Dong, L., Chen, W., Li, W., Xiao, G., Zhao, J., Hao, Y., Xu, Y., Gao, Y., Liu, W., Liu, Y., Yin, H., Li, J., Li, X., Zhao, Y., Wang, X., Ni, F., Ma, X., Li, A., Xu, S.S., Bai, G., Nevo, E., Gao, C., Ohm, H., and Kong, L. (2020). Horizontal gene transfer of Fhb7 from fungus underlies Fusarium head blight resistance in wheat. Science 368.

Wang, J., Na, J.K., Yu, Q., Gschwend, A.R., Han, J., Zeng, F., Aryal, R., VanBuren, R., Murray, J.E., Zhang, W., Navajas-Perez, R., Feltus, F.A., Lemke, C., Tong, E.J., Chen, C., Wai, C.M., Singh, R., Wang, M.L., Min, X.J., Alam, M., Charlesworth, D., Moore, P.H., Jiang, J., Paterson, A.H., and Ming, R. (2012). Sequencing papaya X and Yh chromosomes reveals molecular basis of incipient sex chromosome evolution. Proc Natl Acad Sci U S A 109, 13710–13715.

Wang, Y., Cai, X., Zhang, Y., Hörandl, E., Zhang, Z., and He, L. (2023). The male-heterogametic sex determination system on chromosome 15 of *Salix triandra* and *Salix arbutifolia* reveals ancestral male heterogamety and subsequent turnover events in the genus *Salix*. Heredity (Edinb) 130, 122–134.

Weir, B.S., and Cockerham, C.C. (1984). ESTIMATING F-STATISTICS FOR THE ANALYSIS OF POPULATION STRUCTURE. Evolution 38, 1358–1370.

Westergaard, M. (1958). The mechanism of sex determination in dioecious flowering plants. Adv Genet 9, 217–281.

Wilson Sayres, M.A., and Makova, K.D. (2013). Gene survival and death on the human Y chromosome. Mol Biol Evol 30, 781–787.

Xu, C.Q., Liu, H., Zhou, S.S., Zhang, D.X., Zhao, W., Wang, S., Chen, F., Sun, Y.Q., Nie, S., Jia, K.H., Jiao, S.Q., Zhang, R.G., Yun, Q.Z., Guan, W., Wang, X., Gao, Q., Bennetzen, J.L., Maghuly, F., Porth, I., Van de Peer, Y., Wang, X.R., Ma, Y., and Mao, J.F. (2019a). Genome sequence of *Malania oleifera*, a tree with great value for nervonic acid production. Gigascience 8.

Xu, G.C., Xu, T.J., Zhu, R., Zhang, Y., Li, S.Q., Wang, H.W., and Li, J.T. (2019b). LR_Gapcloser: a tiling path-based gap closer that uses long reads to complete genome assembly. Gigascience 8.

Xu, L., Wa Sin, S.Y., Grayson, P., Edwards, S.V., and Sackton, T.B. (2019c). Evolutionary Dynamics of Sex Chromosomes of *Paleognathous* Birds. Genome Biol Evol 11, 2376–2390.

Xue, L., Wu, H., Chen, Y., Li, X., Hou, J., Lu, J., Wei, S., Dai, X., Olson, M.S., Liu, J., Wang, M., Charlesworth, D., and Yin, T. (2020). Evidences for a role of two Y-specific genes in sex determination in *Populus deltoides*. Nat Commun 11, 5893.

Yang, W., Wang, D., Li, Y., Zhang, Z., Tong, S., Li, M., Zhang, X., Zhang, L., Ren, L., Ma, X., Zhou, R., Sanderson, B.J., Keefover-Ring, K., Yin, T., Smart, L.B., Liu, J., DiFazio, S.P., Olson, M., and Ma, T. (2021). A General Model to Explain Repeated Turnovers of Sex Determination in the Salicaceae. Mol Biol Evol 38, 968–980.

Yang, Z. (2007). PAML 4: phylogenetic analysis by maximum likelihood. Mol Biol Evol 24, 1586–1591.

Yue, J., Krasovec, M., Kazama, Y., Zhang, X., Xie, W., Zhang, S., Xu, X., Kan, B., Ming, R., and Filatov, D.A. (2023). The origin and evolution of sex chromosomes, revealed by sequencing of the *Silene latifolia* female genome. Curr Biol 33, 2504–2514 e2503.

Zhang, C., Rabiee, M., Sayyari, E., and Mirarab, S. (2018). ASTRAL-III: polynomial time species tree reconstruction from partially resolved gene trees. BMC Bioinformatics 19, 153.

Zhang, H., Sigeman, H., and Hansson, B. (2022a). Assessment of phylogenetic approaches to study the timing of recombination cessation on sex chromosomes. J Evol Biol 35, 1721–1733.

Zhang, S., Wu, Z., Ma Zhai, J., Han, X., Jiang, Z., Liu, S., Xu, J., Jiao, P., and Li, Z. (2022b). Chromosome-scale assemblies of the male and female *Populus euphratica* genomes reveal the molecular basis of sex determination and sexual dimorphism. Commun Biol 5, 1186.

Zhang, Z., and Yu, J. (2006). Evaluation of six methods for estimating synonymous and nonsynonymous substitution rates. Genomics Proteomics Bioinformatics 4, 173–181.

Zhou, R., Macaya-Sanz, D., Carlson, C.H., Schmutz, J., Jenkins, J.W., Kudrna, D., Sharma, A., Sandor, L., Shu, S., Barry, K., Tuskan, G.A., Ma, T., Liu, J., Olson, M., Smart, L.B., and DiFazio, S.P. (2020). A willow sex chromosome reveals convergent evolution of complex palindromic repeats. Genome Biol 21, 38.

