## Supplementary figures for "Gap-free X and Y chromosomes of *Salix arbutifolia* reveal an evolutionary change from male to female heterogamety in willows, without a change in the sex-determining region"

**Figure S1**

Chromosome quotients (CQ) in 50-kb nonoverlapping window of haplotype *a* of *S. arbutifolia*.

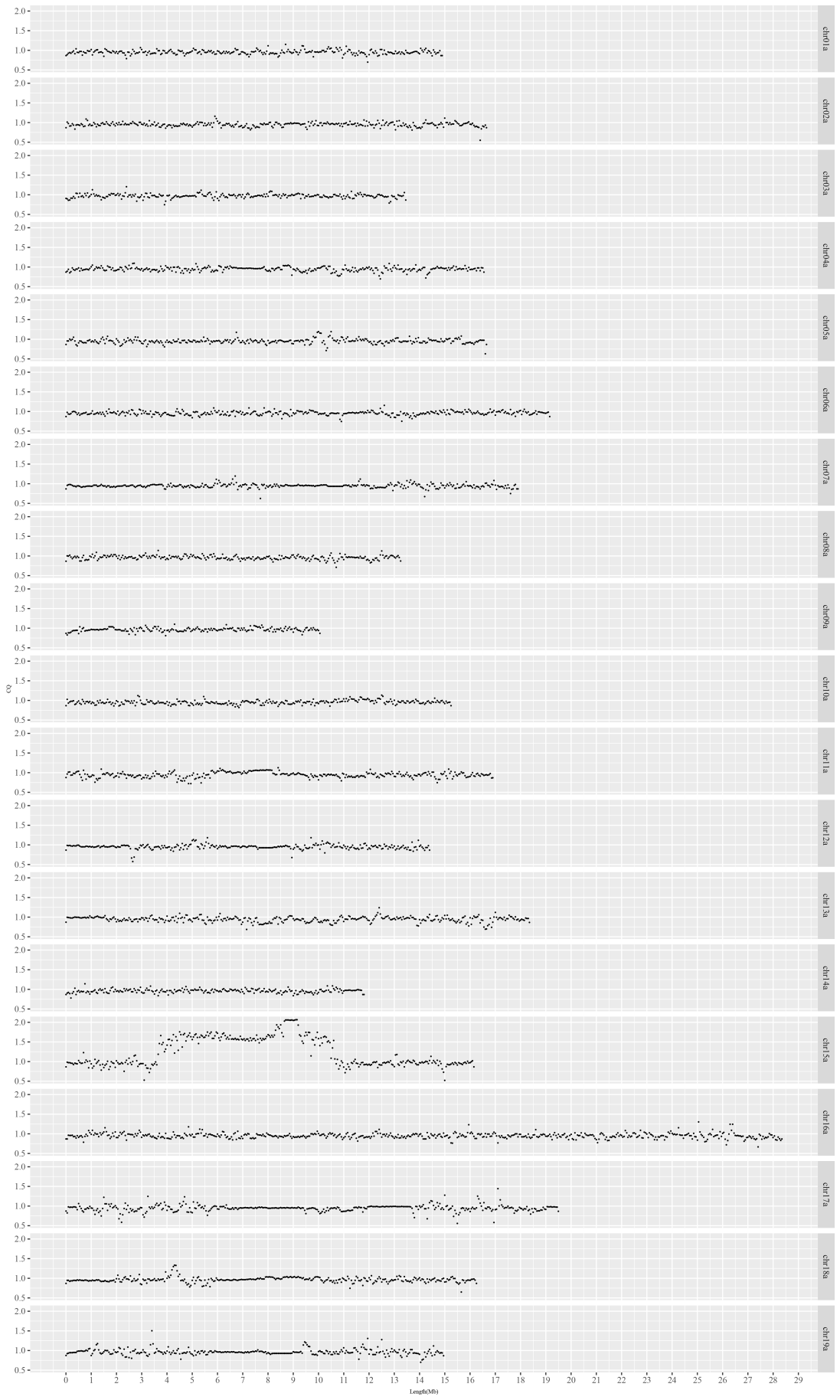

**Figure S2**

Chromosome quotients (CQ) in 50-kb nonoverlapping window of haplotype *b* of *S. arbutifolia*.

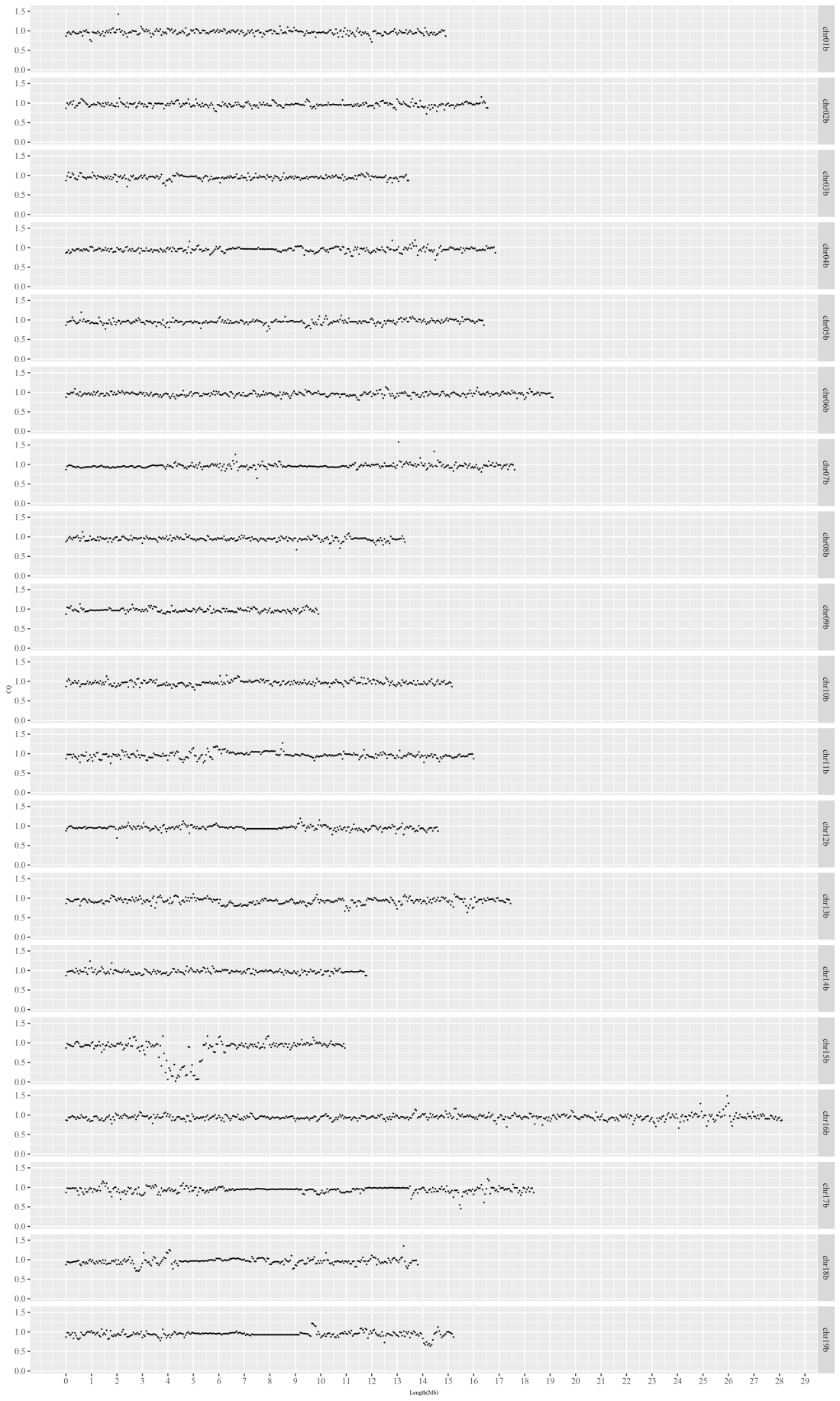

**Figure S3**

F_ST_ values between the sexes for 100-kb overlapping windows of haplotype *a* calculated at 10-kb steps.

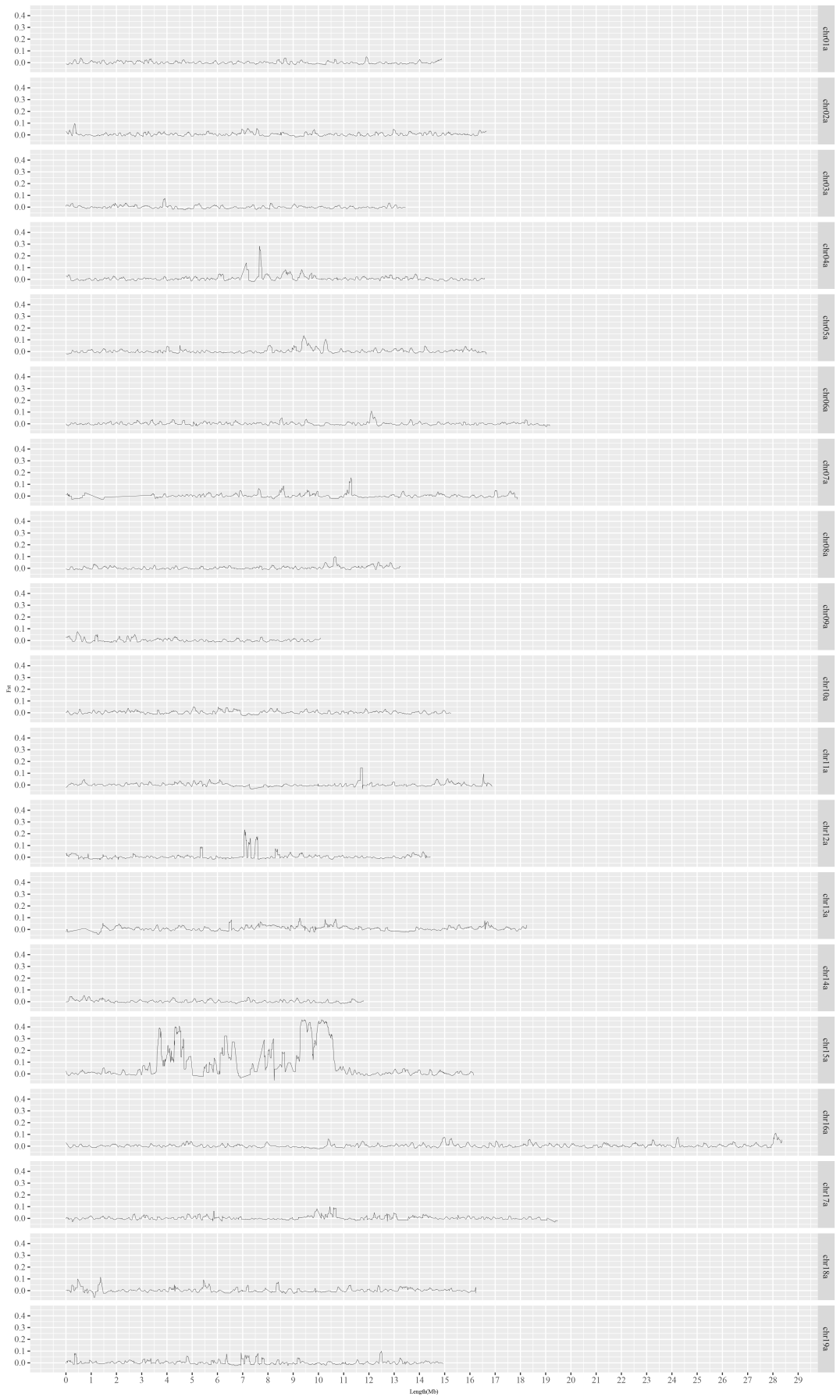

**Figure S4**

F_ST_ values between the sexes for 100-kb overlapping windows of haplotype *b* calculated at 10-kb steps.
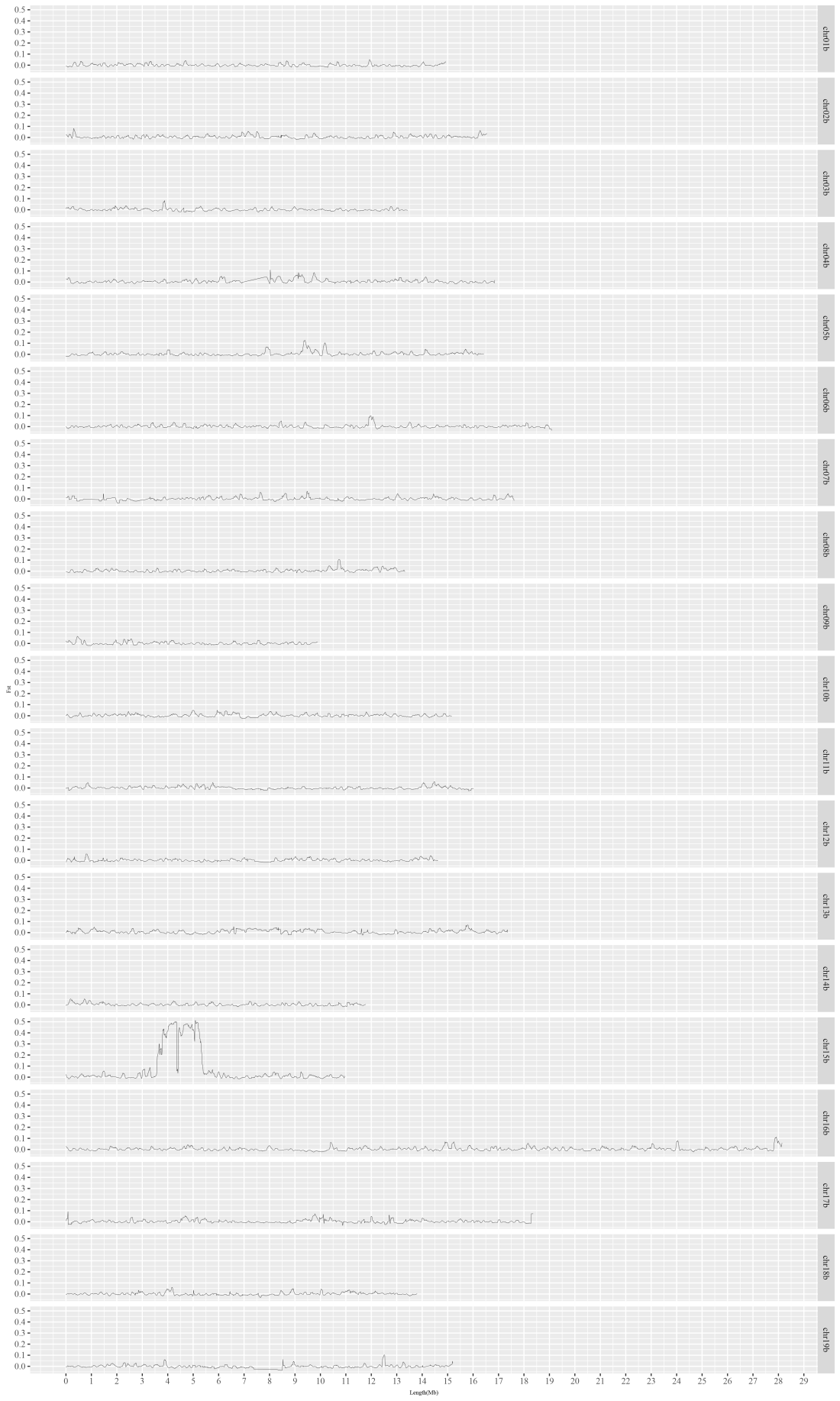

**Figure S5**

(a) Quantile–Quantile (Q–Q) plots of observed and expected GWAS P-values of *Salix arbutifolia* haplotype *a*. (b) Results of GWAS between SNPs and the sexes of 39 individuals using *Salix arbutifolia* haplotype *a* as reference. The y axis is the negative logarithm of *p* values. The red line shows the Bonferroni-corrected significance level corresponding to α < 0.05.

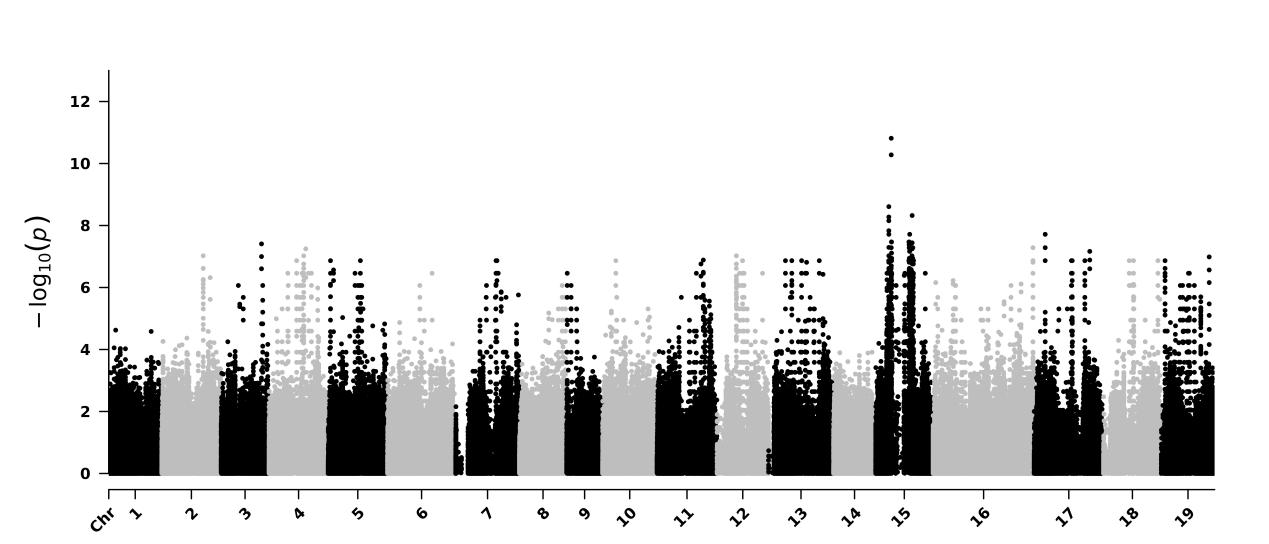

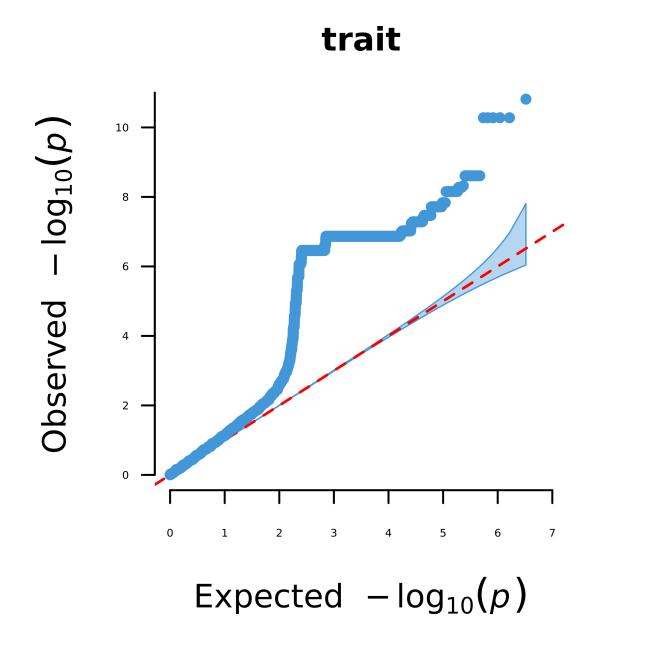

(a)

(b)

**Figure S6**

(a) Quantile–Quantile (Q–Q) plots of observed and expected GWAS *P*-values of *Salix arbutifolia* haplotype *b*. (b) Results of genome wide association studies (GWAS) between SNPs and the sexes of 39 individuals using *Salix arbutifolia* haplotype *b* as reference. The y axis is the negative logarithm of *p* values. The red line shows the Bonferroni-corrected significance level corresponding to α < 0.05.

(a)

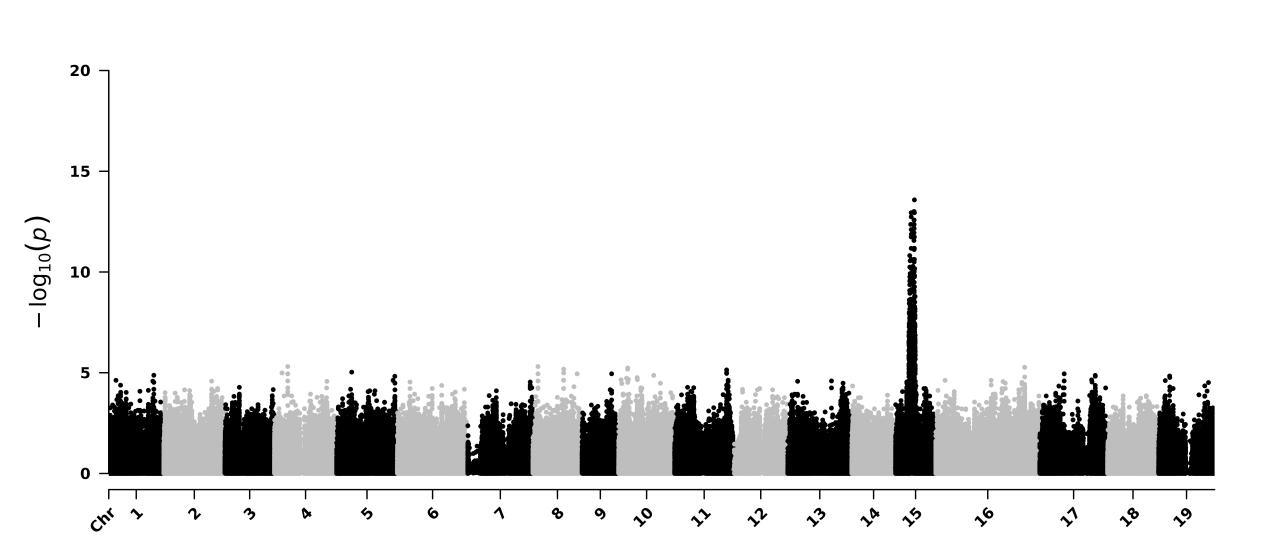

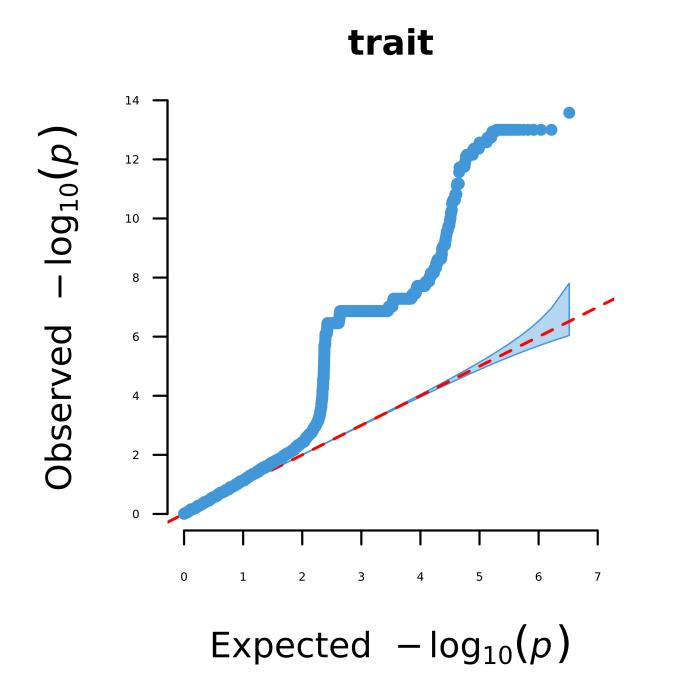

(b)

**Figure S7**

(a) Circos plot of all 38 chromosomes. a, the chromosome lengths in Mb; b, gene density; c, total TE density; d, LTR-Gypsy density and e, LTR-copia density. Blue regions on each chromosome represent putative centromere regions. (b) Magnification of gene density, total TE density, LTR-Gypsy density, LTR-copia density of 15X and 15Y. The red dotted line represents the boundaries of sex-linked regions, the blue region represents the inferred centromeric region, and the green region represents the ancestral sex-linked region (or genes, see below).

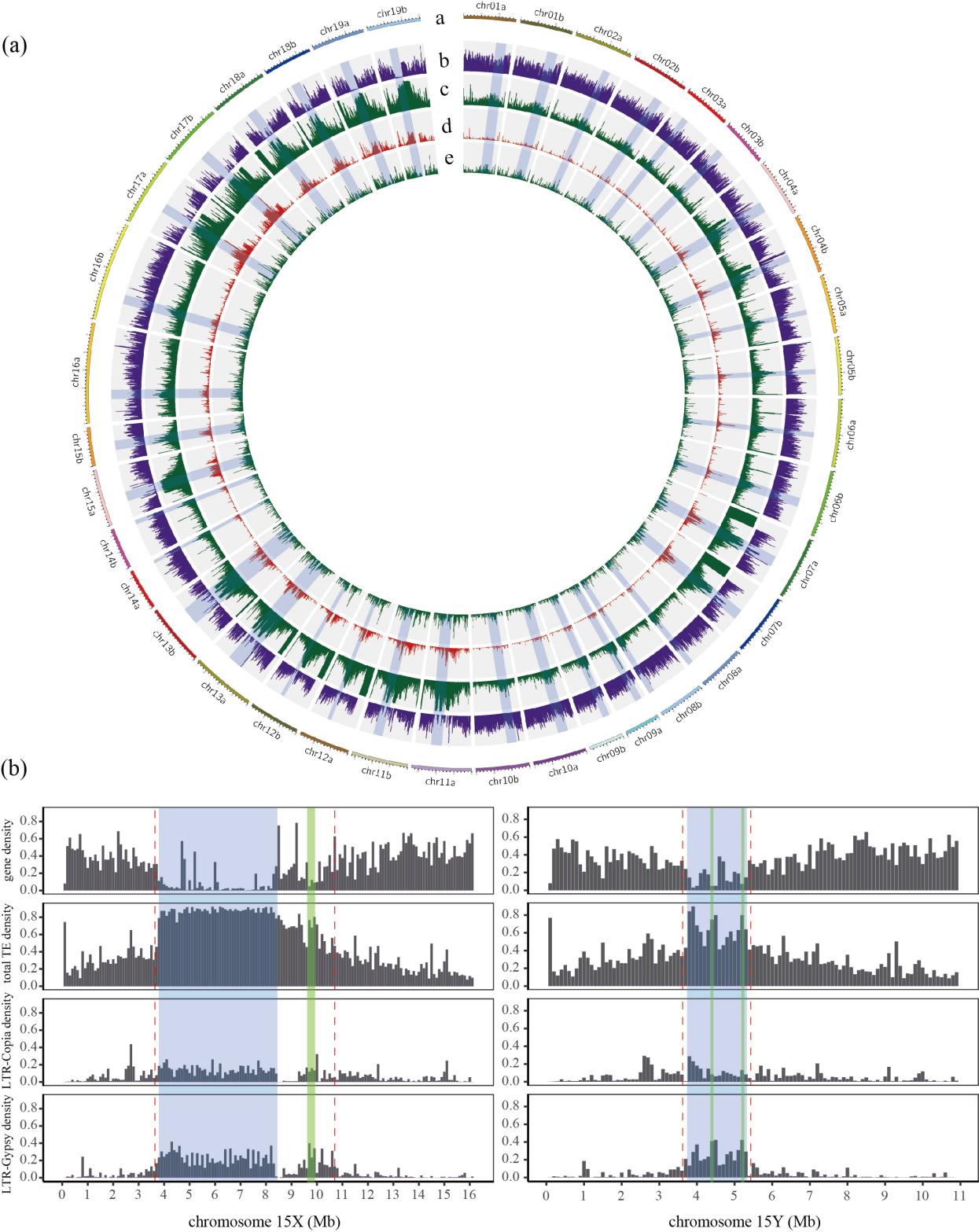

**Figure S8**

Phylogenetic tree constructed using the sequences of both intact and partial ARR17-like genes found in the reference genomes of *Salix* species and of *Populus trichocarpa*. The homologous ARR17-like gene in *Arabidopsis thaliana* was used as the outgroup to all the sequences. The chromosome on which the sequence was found in each *Salix* species is indicated, and the species are indicated as follows: SdunniiF: *Salix dunnii*; Saarb: *Salix arbutifolia*; Spur: *Salix purpurea*; Potr: *Populus trichocarpa.* The chromosome on which the sequence was found in each species is indicated.

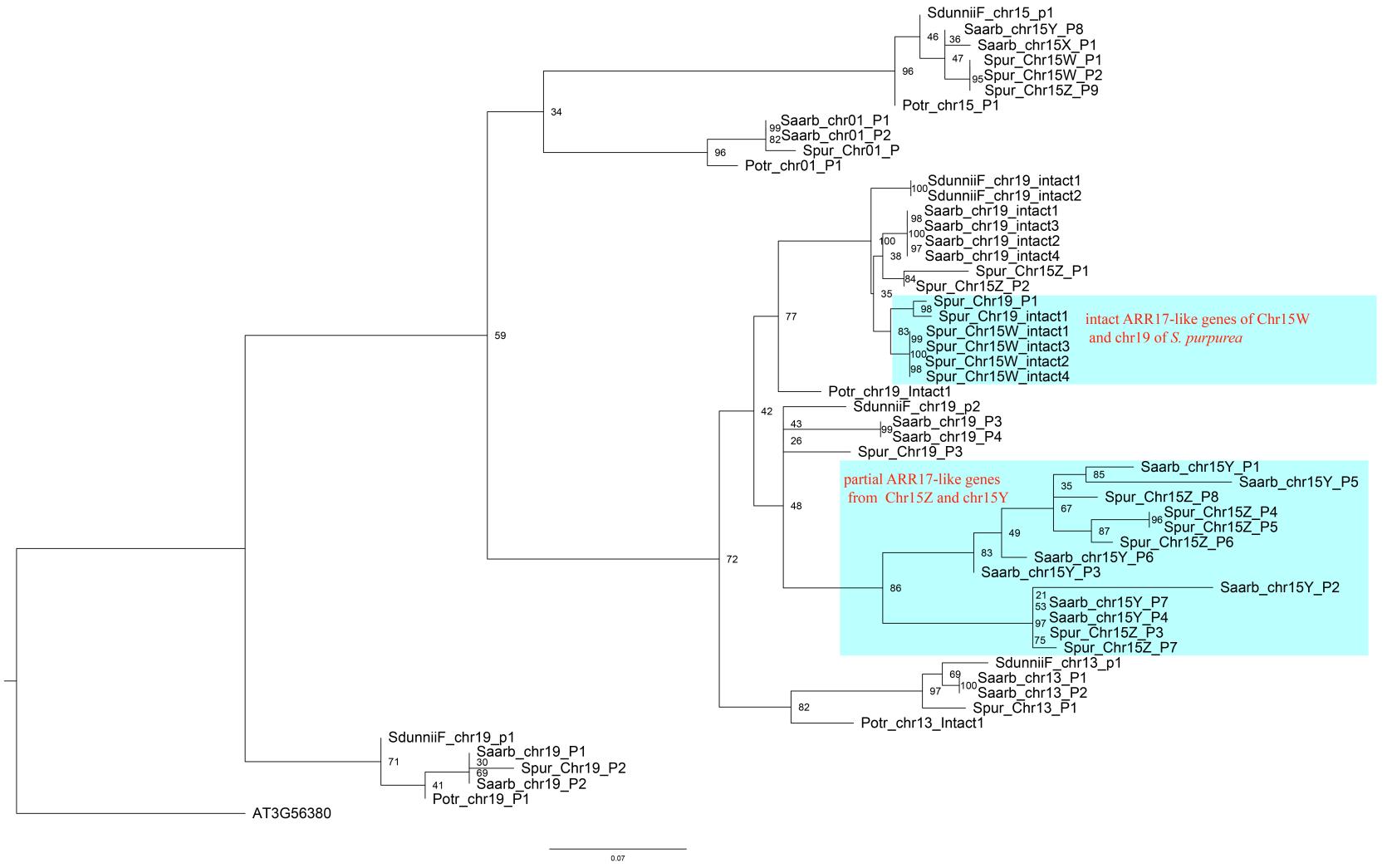

**Figure S9**

Distribution of Ks values between homologous genes in *S. arbutifolia* 15X and 15Y, and *S. purpurea* 15Z-SLR and 15 W-SLR. (a) *S. arbutifolia* 15X and 15Y, the abscissa is the chromosome length (Mb), and the ordinate is the Ks values. Different colored areas represent different evolutionary stratum, and from left to right are putative possible stratum 2, stratum 1, and possible stratum 3. Areas with white color represent PARs. (b) *S. purpurea* 15Z-SLR and 15 W-SLR, the abscissa only shows the phased Z-SLR, the ordinate is the Ks value, and the colored areas are possible stratum 2 and stratum 1 from left to right.

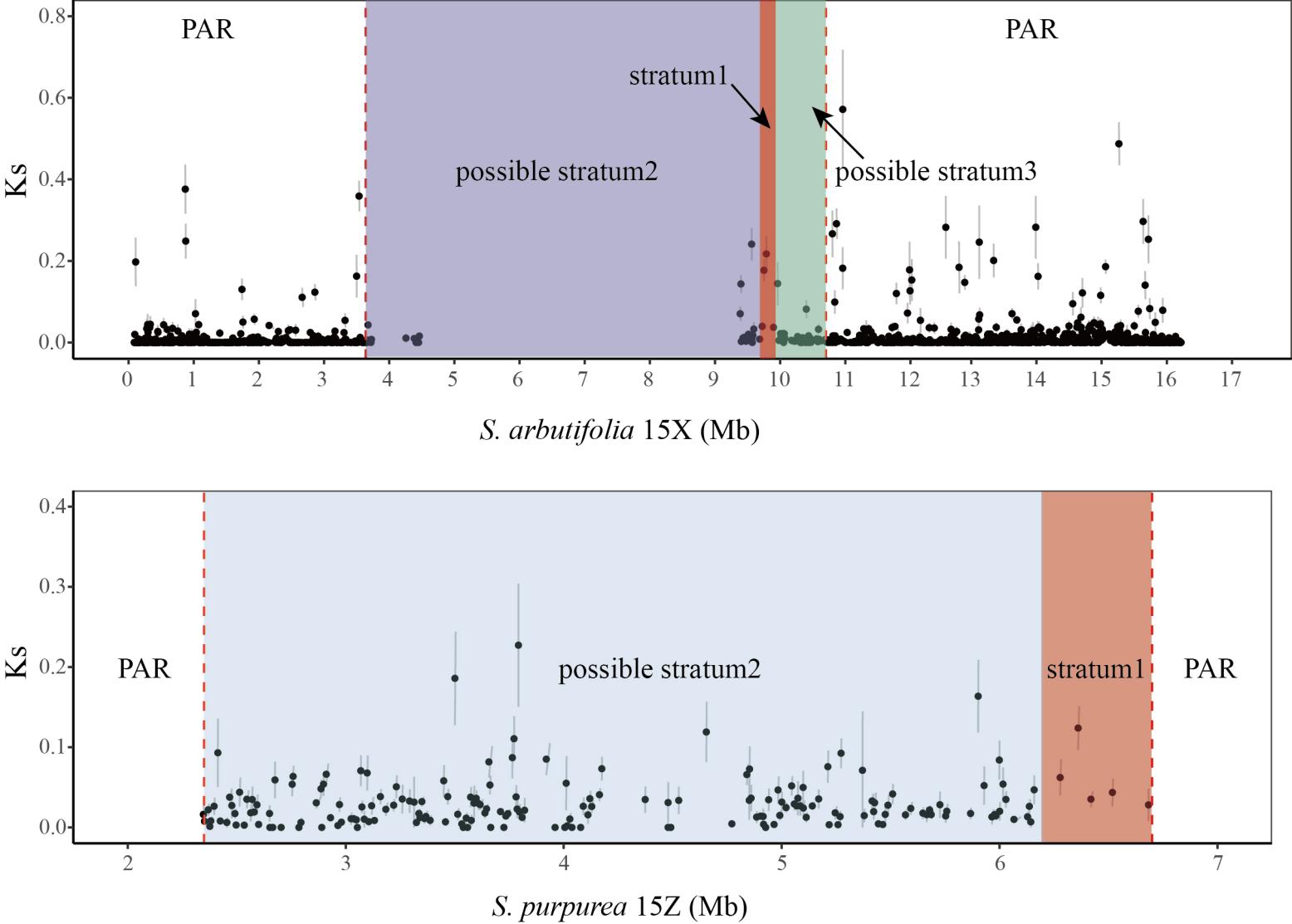

**Figure S10**

Synteny analysis of *S. dunnii* autosome 15 and *S. purpurea* ZW chromosomes. The dark gray region represents the Z and W sex-linked regions and the corresponding collinear region of *S. dunnii* autosome 15.

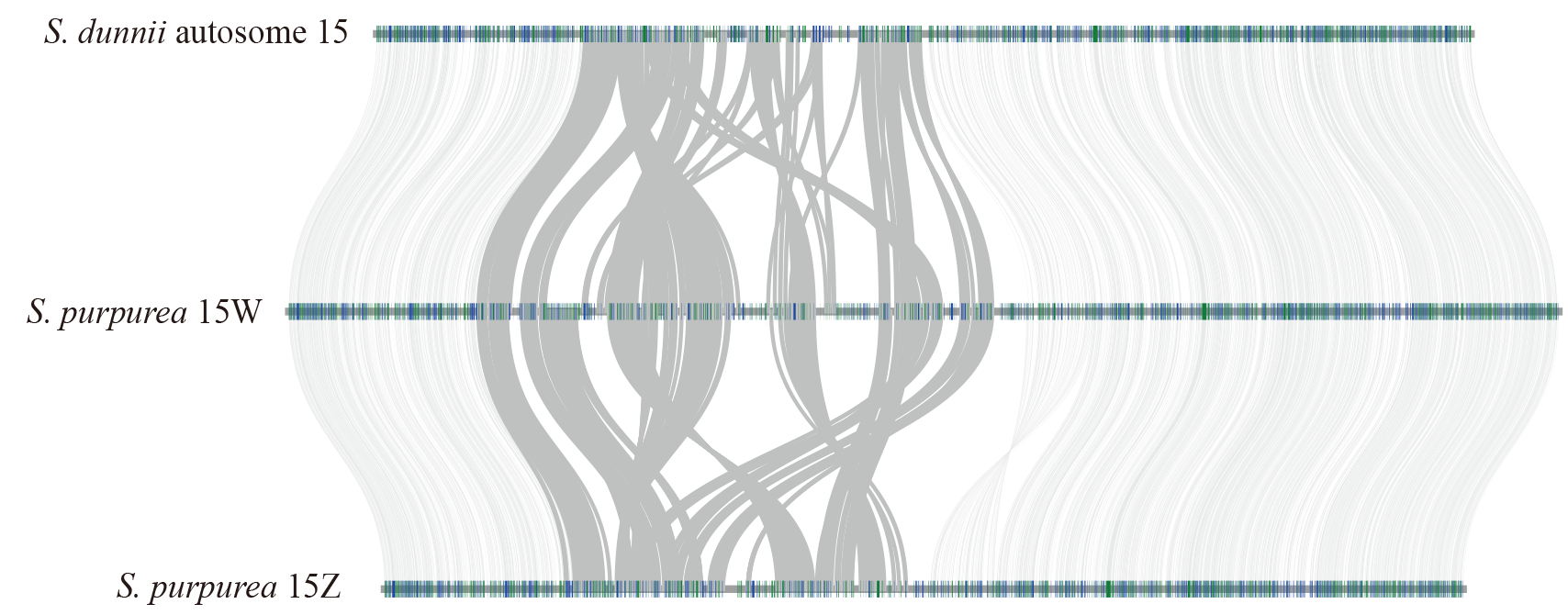

**Figure S11**

Collinearity analysis using *S. purpurea* 15Z, *S. arbutifolia* 15Y and *S. dunnii* autosome 15. The region within the blue line represents the 15Z sex-linked region and the corresponding collinear region of *S. arbutifolia* 15Y and *S. dunnii* autosome 15. The region within the black line represents the *S. arbutifolia* 15Y sex-linked region and the corresponding collinear region of *S. purpurea* 15Z and *S. dunnii* autosome 15.

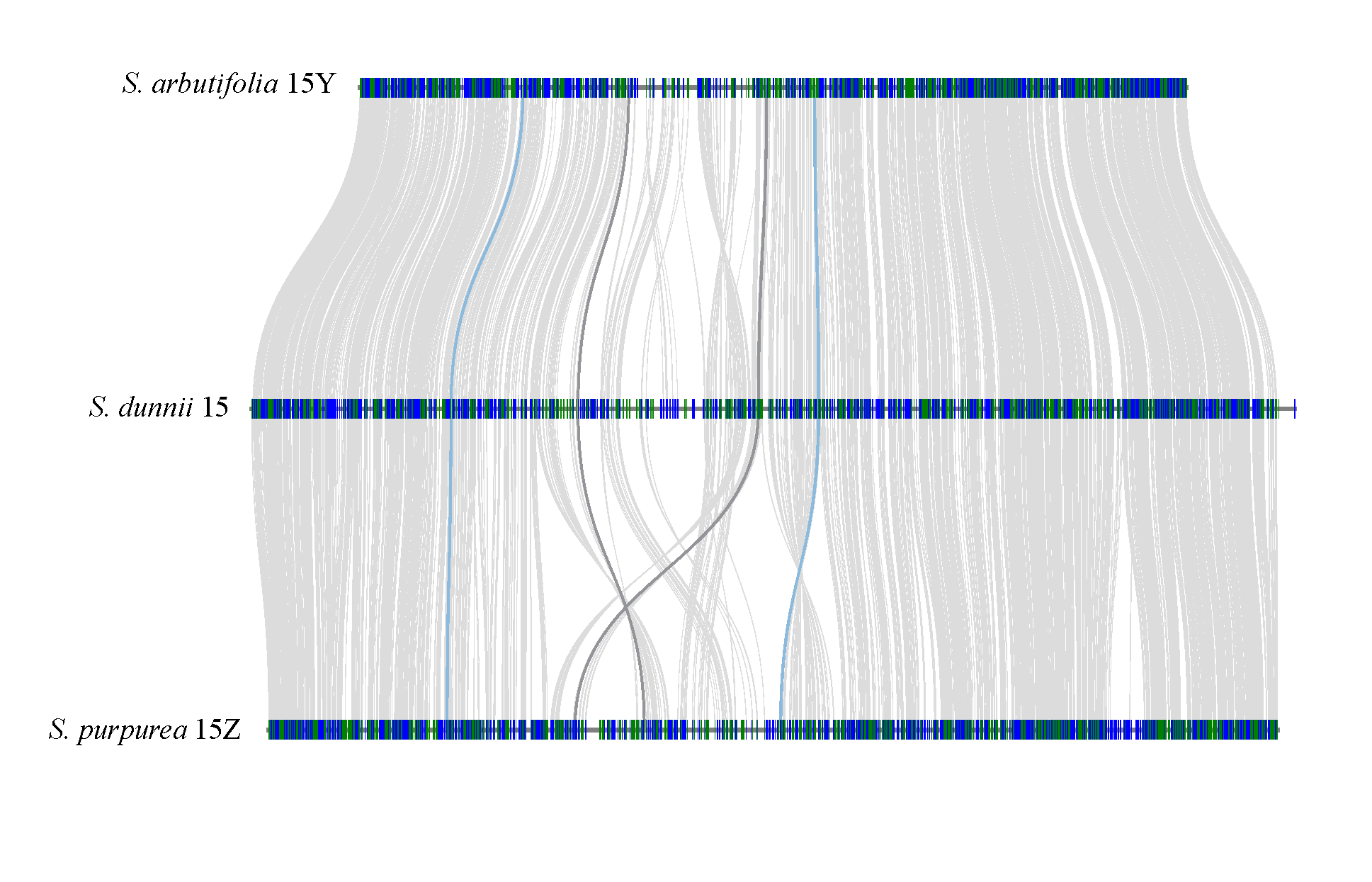
